# Anal skin-like epithelium mediates colonic wound healing

**DOI:** 10.1101/2021.06.02.446836

**Authors:** Cambrian Y. Liu, Nandini Girish, Marie L. Gomez, Philip E. Dubé, M. Kay Washington, Benjamin D. Simons, D. Brent Polk

## Abstract

Intestinal epithelial wound healing, which is essential for health, is compromised and represents a therapeutic target in inflammatory bowel disease (IBD). While studies have elucidated important subpopulations of intestinal epithelial cells in repair, these have yet to translate to therapies. Here, in mouse models of acute colitis, we demonstrate a distinct and essential source of wound-healing cells that re-epithelialize the distal colon. Using 3-d imaging, lineage tracing, and single-cell transcriptomics, we show that neighboring skin-like (squamous) cells of the anus rapidly migrate into the injured colon and establish a permanent epithelium of crypt-like morphology. These squamous cells derive from a small unique transition zone, at the boundary of colonic and anal epithelium, that resists colitis. The cells of this zone have a pre-loaded program of colonic differentiation and further upregulate key aspects of colonic epithelium during repair. Thus, heterologous cell-types at tissue junctions represent unique reserve cells capable of repair and plasticity.

## Introduction

Defects in epithelial barrier function and repair have been linked to chronic inflammation in a variety of organ systems (Elias, 2008; Georas and Rezaee, 2014; Lichtenberger et al., 2013; McCole, 2014). In inflammatory bowel disease (IBD), intestinal epithelial damage accelerates inflammation as part of a vicious cycle. Direct healing of the epithelial barrier is an attractive therapeutic target; however, the cellular programs mediating wound repair remain to be elucidated and leveraged. Wound healing in adult mammals is associated with reprogramming or dedifferentiation of surviving epithelium (Shivdasani et al., 2021), but this generally occurs within a lineage-restricted paradigm, e.g. (Donati et al., 2017; Rinkevich et al., 2011; Tetteh et al., 2016; van Es et al., 2012). Whether the wound healing response in colitis could involve the recruitment of non-intestinal cells remains in question.

During the analysis of three-dimensional, whole-mount reconstructions of murine colonic mucosa during repair from acute colitis, we have observed an unusual skin-like epithelial structure in the rectum (Hale and Cianciolo, 2008; Liu et al., 2015; Seamons et al., 2013). We hypothesized that this structure represents part of the regenerative response of the colon during colitis. Here we define its tissue-of-origin: a molecularly unique transition zone (TZ) of squamous cells in the anus, distal to the anorectal junction. We show that squamous cells are essential for complete colonic re-epithelialization and establish a long-lived epithelium with hybrid colonic-squamous properties as part of the overall repair process. Thus, colitis engages a specialized set of junctional squamous cells, poised for repair, that contribute to the regenerative response and initiate mucosal healing. Our findings define a distinct mode of colonic epithelial repair that originates from neighboring non-colonic skin-like cells.

## Results

### Squamous epithelium contributes to wound healing in colitis

To characterize colonic wound healing processes, we first orally administered 3% dextran sulfate sodium (DSS) to mice for 6 days (d). This model induces acute injury with epithelial erosions (ulcerations) and inflammation, followed by spontaneous healing in the distal colon (rectum) over a span of ~3 weeks (wks) (Liu et al., 2015). In whole-mount total reconstructions of the injured mucosa, obtained with a clearing and imaging protocol we developed previously (Liu et al., 2015), we observed the full extent of a stratified “neo-epithelium” in the distal colon. This neo-epithelium was proximal to the dentate line that, in the uninjured animal, sharply divided KRT8+ colonic (columnar) epithelium from KRT14+ anal (squamous) epithelium (Figure 1a,b, Figure S1a). The neo-epithelium was endoscopically visible (Figure S1b). Microscopically the neo-epithelium resembled thick squamous (skin-like) tissue, but lacked a cornified outer layer (Figure 1c,d). We thus refer to this tissue as <u>s</u>quamous <u>n</u>eo-<u>e</u>pithelium of <u>c</u>olon (SNEC). The KRT14+ SNEC was present from the end of the DSS treatment (exp d 6) and persisted for life, exhibiting basal squamous expression of COL17A1 and the proliferative marker KI-67 (longest timepoint: 15 months post-DSS, Figure 1e, Figure S1c). SNEC grew with repeated cycles of DSS (Figure 1f). We have observed SNEC to occupy up to 1 cm of distal mouse colon (equivalent ~15% of the overall colonic length or ~50% of the rectum) after injury. Patches of SNEC were always contiguous with the dentate line. Three-dimensional reconstructions revealed a field of filled intestinal crypt-like invaginations (Figure 1g), which we termed rete pegs based on their similarity to structures found in skin and mucous membranes. Rete pegs of SNEC, as well as basal cells of anal squamous epithelium next to the dentate line, stained positively for the progenitor cell markers (Arnold et al., 2011; Pellegrini et al., 2001) P63 and SOX2. P63 and SOX2 were not expressed in colonic epithelium before or after DSS treatment (Figure 1h). Thus, acute injury and inflammation of the distal colon are associated with the formation of a permanent squamous tissue structure.

**Figure 1:**
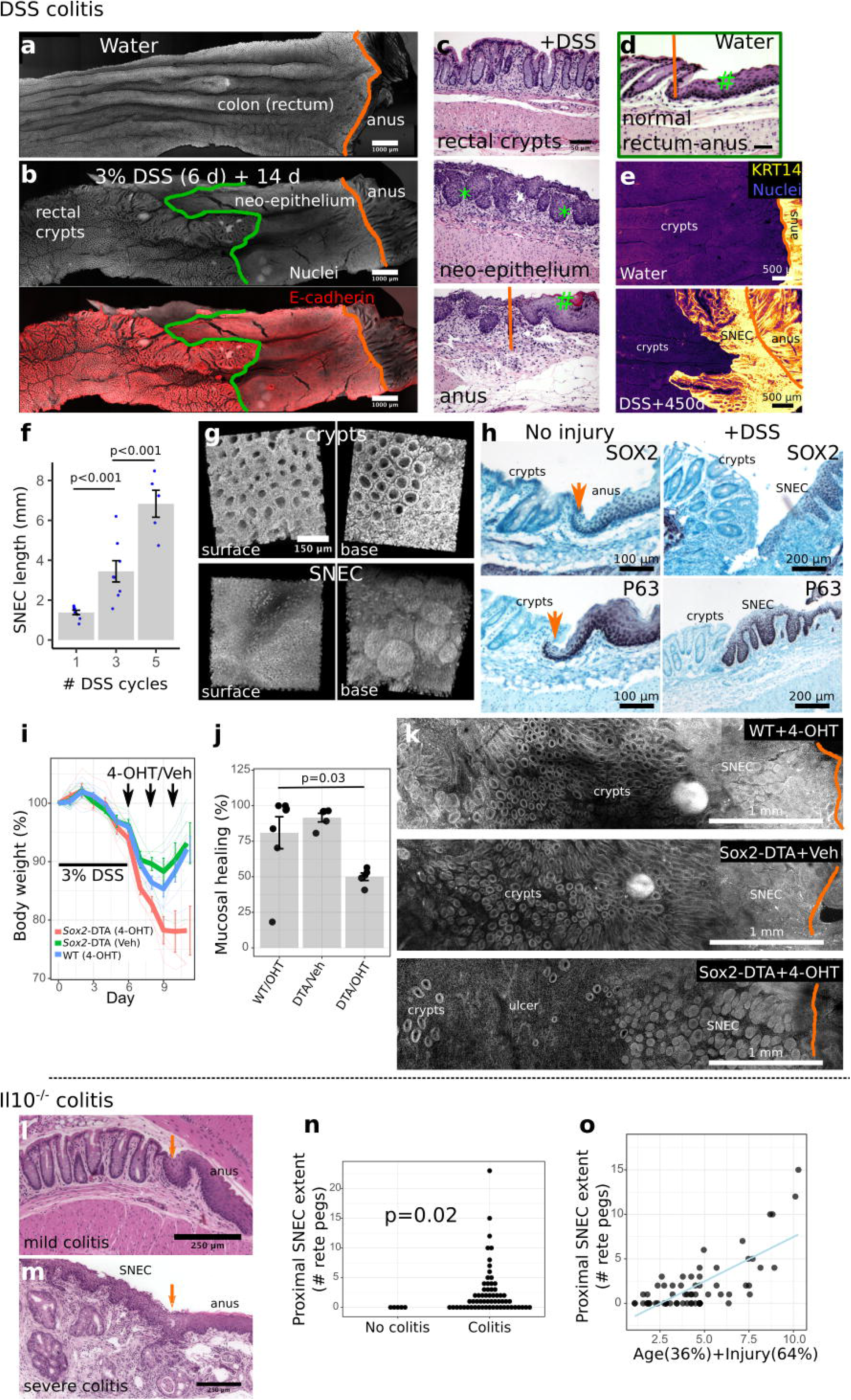
Squamous epithelium contributes to wound healing in colitis. a-b) Surface representations after 3D imaging of chemically cleared, flattened colonic tissue after DSS injury reveal a distinct epithelium replacing colonic crypts, proximal to the dentate line (orange) defining the sharp anorectal epithelial transition in uninjured specimens, but distal to the margin (green) of colonic crypts. c-d) H&E images show that this long-lasting “neo-epithelium” (c) has a broadly squamous structure with rete pegs (*) but lacks a fully differentiated surface (#). The neo-epithelium does not exist in the absence of injury (d). e) Squamous neo-epithelium of colon (SNEC) persists as a KRT14+ tissue structure 15 months after injury (n=3 mice), as shown in whole-mount images. f) Plot of SNEC size (length of incursion from the dentate line) with multiple cycles of DSS inducing repetitive injury (n=5-7 mice/condition). g) 3D reconstruction of SNEC surface and abluminal view shows distinct morphology compared to colon crypts. h) Immunostaining reveals high expression of P63 and SOX2 in squamous cells at the dentate line and in rete pegs, but exclusion from colonic epithelium (n=4 mice). Orange arrows: dentate line. i-k) Anorectal squamous cell apoptosis was induced by genetic targeting of intracellular diphtheria toxin protein A (DTA) to Sox2+ cells (Sox2-DTA mice), with conditional expression induced by tamoxifen enemas (4-OHT). Apoptotic targeting retards body weight recovery (i), leaving ulcers (j,k) in the colon after DSS colitis (n=5 mice/condition). Significance was evaluated using the t-test with false discovery rate correction. l) Retrospective analysis of squamous epithelium in colons of Il10-/-mice. H&E photomicrographs show that young (8 wks-old) Il10-/-mice lacking inflammation were devoid of SNEC. Orange arrow marks the dentate line. m) However, older mice (>36 wks-old) with severe distal colitis exhibit SNEC. n) Colitis (as defined by histological injury score >0) was a prerequisite for finding SNEC. Each dot represents a mouse. o) A combination of age and histological colitis score modeled the size of the SNEC tissue, in agreement with SNEC being prevalent in older animals with extensive illness. Error bars: SE. See also Figures S1-S3.

The ability of SNEC to rapidly re-epithelialize the colonic surface suggests that it plays a functional role in wound healing. To define the functional contribution of anal-derived cells to outcomes of colonic injury, we targeted apoptotic signals to Sox2+ cells at the anorectal junction. In whole-tissue (Figure S2a) and single-cell cell transcriptomic (Figure S2b) datasets described below and analyzed retrospectively, the expression of Sox2 in squamous tissues was >10-fold higher than in the colon, which was confirmed with immunohistochemistry (Figure 1h). Sox2 expression remained low in colon samples during DSS injury. Thus, targeting of Sox2+ cells would largely spare colonic epithelium. We generated Sox2::CreER (Arnold et al., 2011);Rosa26::LSL-DTA (Voehringer et al., 2008) (Sox2-DTA) mice, which activate in response to tamoxifen the intracellular expression of the diphtheria toxin A (DTA) polypeptide chain in Sox2-expressing cells. Enema administration of low doses of the topically active tamoxifen metabolite 4-OHT induced specific apoptosis after 2 d in anal epithelium, but not in other alimentary organs (Figure S2c). Delivery of 4-OHT during the acute recovery period after DSS exposure prevented the normal rebound in body weight (Figure 1i) and left residual ulcers (Figure 1j), reflective of impaired mucosal healing (Figure 1k), as assessed from 3d reconstructions. Intriguingly, squamous cells resembling SNEC and anal epidermis remained detectable adjacent to the residual ulcers and may have partially escaped ablation or undergone rapid regeneration. To determine if these outcomes were specific to the period of acute injury, we targeted apoptosis to SNEC cells 4 wks after injury. Although tamoxifen induced apoptosis in SNEC (Figure S2d), enema administration of 4-OHT at this late timepoint did not affect subsequent body weights (Figure S2e). Thus, apoptotic targeting to anorectal squamous cells during acute injury impedes both ulcer repair and clinical recovery from colitis.

To determine if chronic colitis was also associated with formation of SNEC, we studied archived colonic samples (Dube et al., 2012) obtained from Il10-/-mice, which spontaneously develop a progressive colitis (Kuhn et al., 1993). We observed SNEC in animals with severe colitis (Figure 1l,m). Consistent with results observed with DSS-induced colitis, we did not find SNEC in uninjured specimens (Figure 1n). To quantify the relationship between SNEC size and injury, we plotted the relationship between rete peg number and age and histological score. A linear mixture of age and histological score predicted the extent of SNEC (Figure 1o). Thus, in a chronic colitis model, the duration and severity of injury both predicted the formation of SNEC.

The ontogeny of SNEC may be related to the response to tissue injury. Moreover, the detection of SNEC in both acute colitis (DSS) and chronic colitis (Il10-/-) models suggested this tissue’s emergence within an inflammatory milieu. To characterize the tissue context of SNEC, we performed whole-mucosal transcriptome profiling of SNEC, colon, and anus during the DSS-induced injury-repair sequence. In a principal component plot (Figure S3a), SNEC samples were initially interposed (exp d 9) between anal and distal colonic samples. However, by exp d 20, SNEC samples more-resembled colonic samples. SNEC samples were marked by the expression of Ifit3 and Ifit3b (Figure S3b), interferon-responsive genes, and were overall associated with immune signaling (Figure S3c). A pathway analysis across all timepoints and tissues demonstrated the onset of inflammatory signaling by exp d 3 in the anus and exp d 6 in the distal colon. While the colon remained largely devoid of proliferative signaling through exp d 6, proliferative signaling persisted at all tested timepoints in anus and SNEC (Figure S3d). Bulk SNEC tissue exhibited remodeling, from a hybrid anorectal phenotype at exp d 9 (“early”), towards the higher expression of intestinal markers at exp d 20 (“late”). This could be due to intermingling of SNEC tissue with differentiated columnar tissue (Figure S3e,f). Thus, SNEC arises within an inflammatory microenvironment, and its constituent squamous cells exhibit persistent and selective proliferative potential throughout the DSS injury cycle.

### SNEC derives from cells of the anal transition zone

We next investigated the tissue origins of SNEC. To determine whether SNEC represented a unique epithelium, we screened for keratin expression in our transcriptomic dataset. We found enriched expression of Krt7 in SNEC (Figure 2a). Prior to injury, a thin transition zone (TZ) of squamous Krt7+ cells, representing a narrow (~10 cell-wide) portion of the anal epithelium, was identified next to the dentate line (Figure 2b). These pre-injury TZ cells and their related post-injury Krt7+ SNEC cells were both negative for markers of anal/skin terminal differentiation (LOR, IVL, KRT1), suggesting that these two tissues are related (Figure 2c, Figure S4).

**Figure 2:**
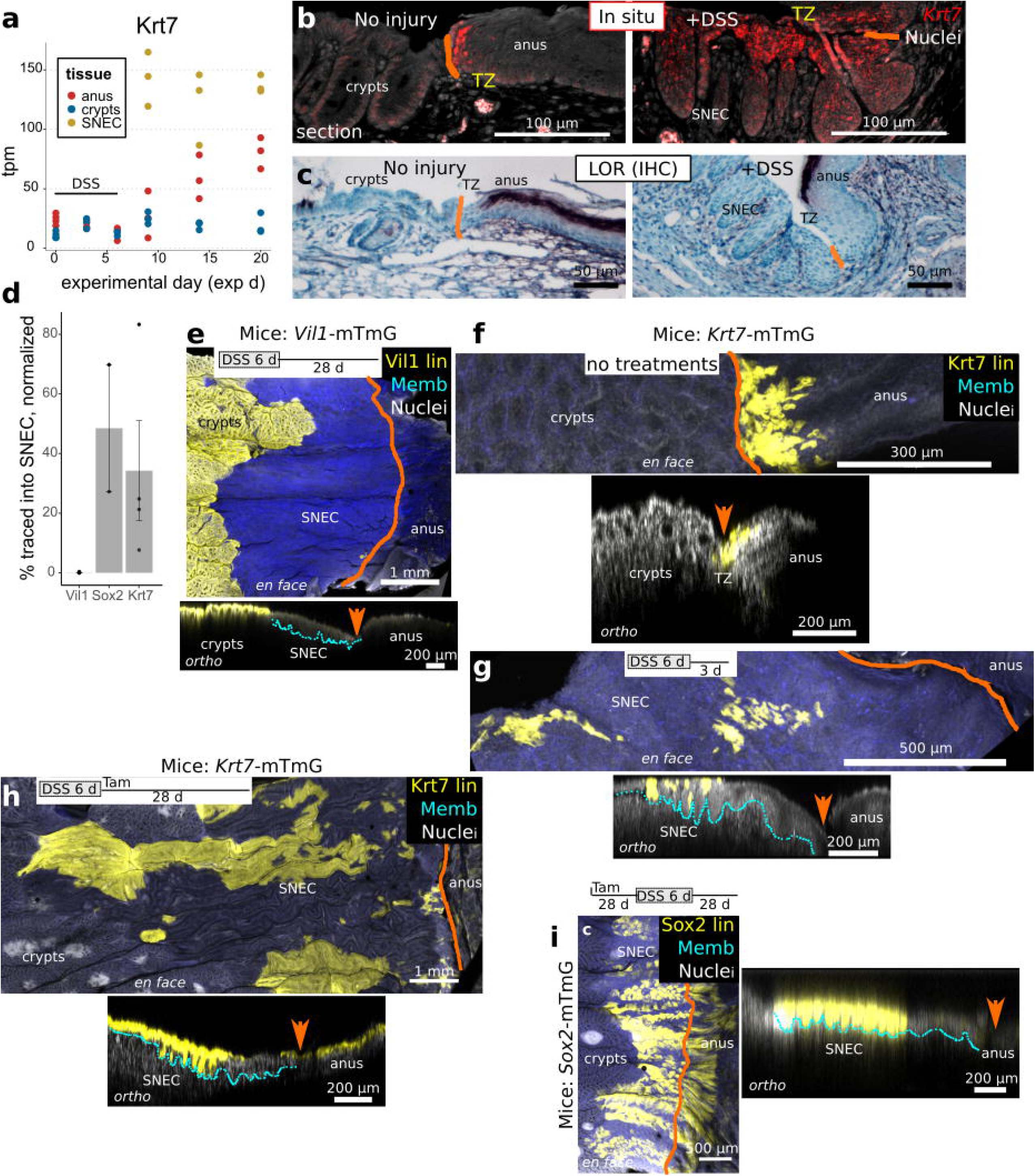
SNEC derives from cells of the anal transition zone. a) Screening of keratin expression in RNA-Seq data from SNEC identifies enrichment of Krt7 (n=3 mice/timepoint). b) In situ hybridization experiments (n=4 mice) demonstrate mRNA expression of Krt7 in SNEC and in a thin transition zone (TZ) of anal squamous cells next to the dentate line (drawn in orange). c) The TZ and SNEC do not express loricrin (LOR), a marker of terminal differentiation of skin (n=5 mice). d) Summary of contributions of various Cre/CreER driver mice to labeling of SNEC; the Vil1+ lineage (n=3 mice) does not contribute whereas Sox2+ (n=2 mice) and Krt7+ (n=4 mice) lineages do. Values represent % of SNEC surface labeled relative to the efficiency of baseline labeling in the colon/TZ/anus. e-i) Visual representations of transgenic labeling data in d, shown as maximum intensity projections of the distal colonic surface (en face) and as rotations (ortho) showing the orthogonal sectioning depth. Crypts were fully labeled in Vil1-mTmG mice, but no labeling was observed in SNEC (e). Outcomes of tracing in Krt7-mTmG mice were assessed in both tamoxifen-free/spontaneous labeling (f,g) (n=4 mice) and 2 mg tamoxifen (n=4 mice) injected-conditions (h). Labeling of TZ and SNEC was observed with both treatment paradigms. The lineages of Sox2+ cells were found in the anal epidermis and in SNEC (i). Error bars: SE. See also Figures S4, S5.

We considered two possibilities for SNEC origins. Colonocytes may have transdifferentiated to squamous cells to produce a TZ-like epithelium. Alternatively, TZ cells migrated into the colon to make SNEC. To distinguish these possibilities, we performed lineage tracing. We compared the representations of Vil1+ (colonic, VIl1-mTmG mice), Sox2+ (anal TZ and anal epidermis, Sox2-mTmG), and Krt7+ (anal TZ, Krt7-mTmG mice) lineages in SNEC (Arnold et al., 2011; el Marjou et al., 2004; Jiang et al., 2017). In homeostasis, these lineage markers labeled the expected tissue (Figure S5). In Krt7-mTmG mice, we observed highly specific labeling of anal TZ cells in the absence of tamoxifen (Figure S5c). After DSS-induced injury, the lineage of Vil1+ cells was not observed in SNEC (Figure 2d,e). Krt7+ cell lineages labeled SNEC across a range of tamoxifen dosages (Figure 2f-h, Figure S5f). We found abundant labeling of SNEC in tamoxifen-treated Sox2-mTmG mice (Fig 2i). The labeling of SNEC included cells near the luminal surface and within the rete pegs (Figure 2f-i). Together, these results suggest that SNEC is derived from Krt7+ Sox2+ anal TZ epithelium and not from colonic crypts.

### Rete pegs progressively form during squamous cell migration

We next assessed the mechanism of rete peg formation within SNEC. In Sox2-Confetti mice, which enable clonal labeling of cells, initiation of tracing at exp d 6 revealed lateral streams of cells that elongated over time, consistent with the proliferation of Sox2+ cells during SNEC formation (Figure S6). We considered that the crypt-like structures of rete pegs might be formed from the passive repopulation of denuded colonic crypts by squamous cells. If true, then rete pegs should be apparent from the beginning of SNEC formation. Alternatively, rete pegs may be formed through an active process within the epithelium, in which a flat surface is gradually compartmentalized. In this case, the appearance of rete pegs would be delayed from the onset of injury.

To differentiate these hypotheses, we examined the dynamics of SNEC formation during the acute phase of DSS-induced colitis and linked them to proliferative changes. We injected mice with EdU and BrdU at 24 h and 2 h, respectively, prior to euthanasia (Figure 3a). The EdU pulse enables fate-tracking of recently proliferative cells. The BrdU pulse locates the active proliferative domain. During the DSS injury-repair cycle, the proliferative potential of colonic epithelium exhibited a near-total loss followed by recovery. The anal TZ and SNEC maintained proliferation throughout (Figure 3b). Prior to injury, EdU+ and BrdU+ cells intermingled within the basal layers of the TZ and anal epidermis. Neighboring rectal epithelium harbored proliferative cells at the basal halves of crypts (Figure 3c). At exp d 6, colonic crypts were devoid of label (Figure 3d). However, at the dentate line, a squamous epithelium lacking rete pegs was found adjacent to the ulceration (Figure 3d’). This flat epithelium was highly proliferative at both exp d 6 and d 8 (Figure 3e). We refer to this expanding front of squamous cells as the “leading edge” of SNEC. By exp d 10, SNEC appeared spatially dichotomized, with a highly proliferative leading edge and a rear (distal) end consisting of rete pegs (Figure 3f). At exp d 43, SNEC was structured exclusively as rete pegs, with proliferative cells exclusively located along the periphery of the pegs (Figure 3g). Thus, SNEC emerges from the anal TZ as a flat epithelium and later acquires the crypt-like rete peg morphology. Consistent with this idea, lateral clonal streams of cells linking SNEC to the anus were initially observed as flat epithelium, and then later observed to be composed of rete pegs (Figure S7).

**Figure 3:**
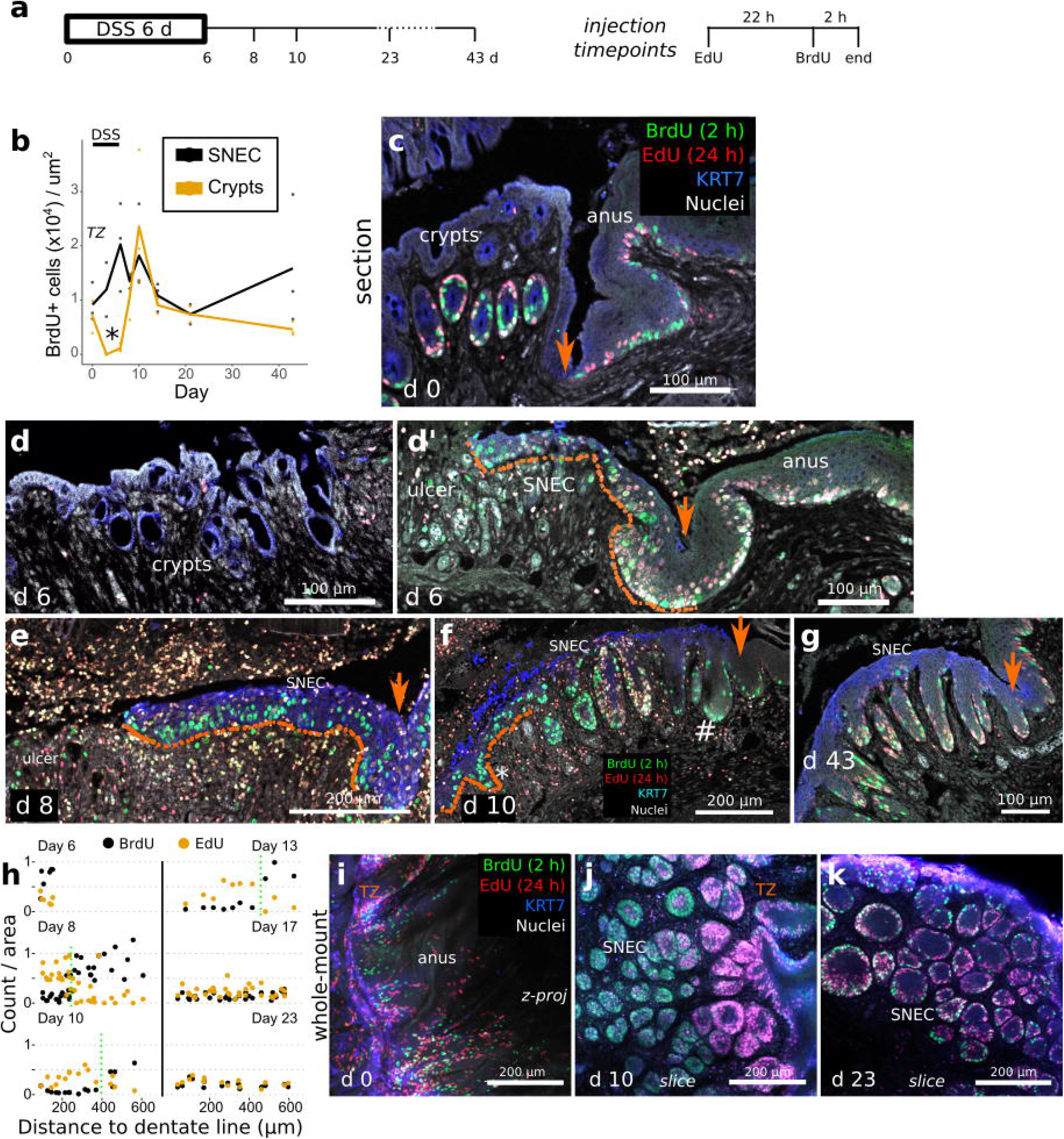
Rete pegs progressively form during squamous cell migration. a) Shown are results of experiments in which mice were injected with 2.5 mg EdU 24 h and 2.5 mg BrdU 2 h prior to dissection, at different timepoints during the DSS injury-repair cycle (n=3 mice/timepoint). Orange arrows mark the dentate line in images. b) Quantification of squamous cell BrdU incorporation vs. colonic crypt incorporation (n=3 mice/timepoint) demonstrates a critical time (exp d 6, asterisk) when SNEC proliferates but colonic epithelium does not. c) In thin sections, at exp d 0, proliferative cells (BrdU+) and their short-term descendants (EdU+) are found in colonic crypts and in the anal TZ and skin. d,d’) At exp d 6, crypts (d) are devoid of proliferative signals, whereas the flat leading edge (dotted line) of SNEC (d’) harbors BrdU+ cells close to the ulcer. e) The flat leading edge (dotted line) of SNEC further expands into the ulcerated region at exp d 8. f) Thin section photomicrograph from at exp d 10, demonstrating new BrdU+ rete pegs (asterisk) forming within the leading edge and a population of proliferation-reduced rete pegs (#) behind them on the right near the dentate line. g) Shown is the reduced nucleotide incorporation at exp d 43 in rete pegs, with label restricted to the periphery. h-k) Quantification of labeled nucleotide incorporation relative to the dentate line (h), from whole-mount images in i-k (n=3 mice/timepoint). The transition between EdU-enriched and BrdU-enriched rete pegs is manually denoted with the dotted green line. i) The homeostatic TZ is a proliferative zone. j) At exp d 10, rete pegs close to the leading edge (left side) are full of BrdU+ cells, indicating rapid proliferation associated with rete peg morphogenesis. However, cells in the interior of rete pegs near the TZ were singly EdU+, suggesting that downregulation of the proliferative rate is involved in maturation of rete pegs. k) Rete pegs at exp d 23 exhibited fewer nucleotide-incorporated cells. See also Figures S6, S7.

To determine how rete peg formation associated with changes in proliferation, we analyzed EdU/BrdU double-labeling experiments in 3d. Quantification of whole-mount images (Figure 3h) showed a highly proliferative (BrdU+) squamous cell population emerging from the KRT7+ TZ at exp d 6 (Figure 3i). However, at exp d 10, there was a proximal-to-distal gradient of labels (Figure 3j). Rete pegs located within proximal regions of SNEC were filled with BrdU+ cells. In contrast, rete pegs located distally and adjacent to the dentate line were filled with exclusively EdU+ cells. This “transition point,” where the characteristic of rete pegs changes from being primarily BrdU+ to primarily EdU+, moved proximally with time (Figure 3h). At exp d 23, all rete pegs exhibited reduced proliferation, with labels restricted to the periphery (Figure 3k). In total, these results are consistent with a simple model in which rete pegs are actively formed from highly proliferative cells within the leading edge. After formation and further distancing from the leading edge, pegs undergo downregulation of proliferation.

### Anal TZ is a molecularly distinct squamous tissue with colonic expression

We next asked what molecular properties of anal TZ were associated with its capability to form a wound-associated squamous tissue in colon. To characterize TZ-derived cells in relation to neighboring epithelia, we performed single-cell RNA-Seq analysis of ~35,000 cells. We obtained these cells from gross dissections of the anorectal junction at 3 different timepoints (exp d 0, 7, and 42) around a DSS injury stimulus (exp d 0-6). These samples contained epithelial and stromal cells of the uninjured TZ, of acutely forming SNEC, and of long-lived SNEC, respectively, in addition to colonocytes and anal epidermal cells recovered from the dissection margins. Dimensional reduction, clustering, and plotting of individual cell types showed transcriptional changes associated with acute injury in SNEC and in neighboring colonic epithelium and anal skin (Figure 4a, Figure S8a-c).

**Figure 4:**
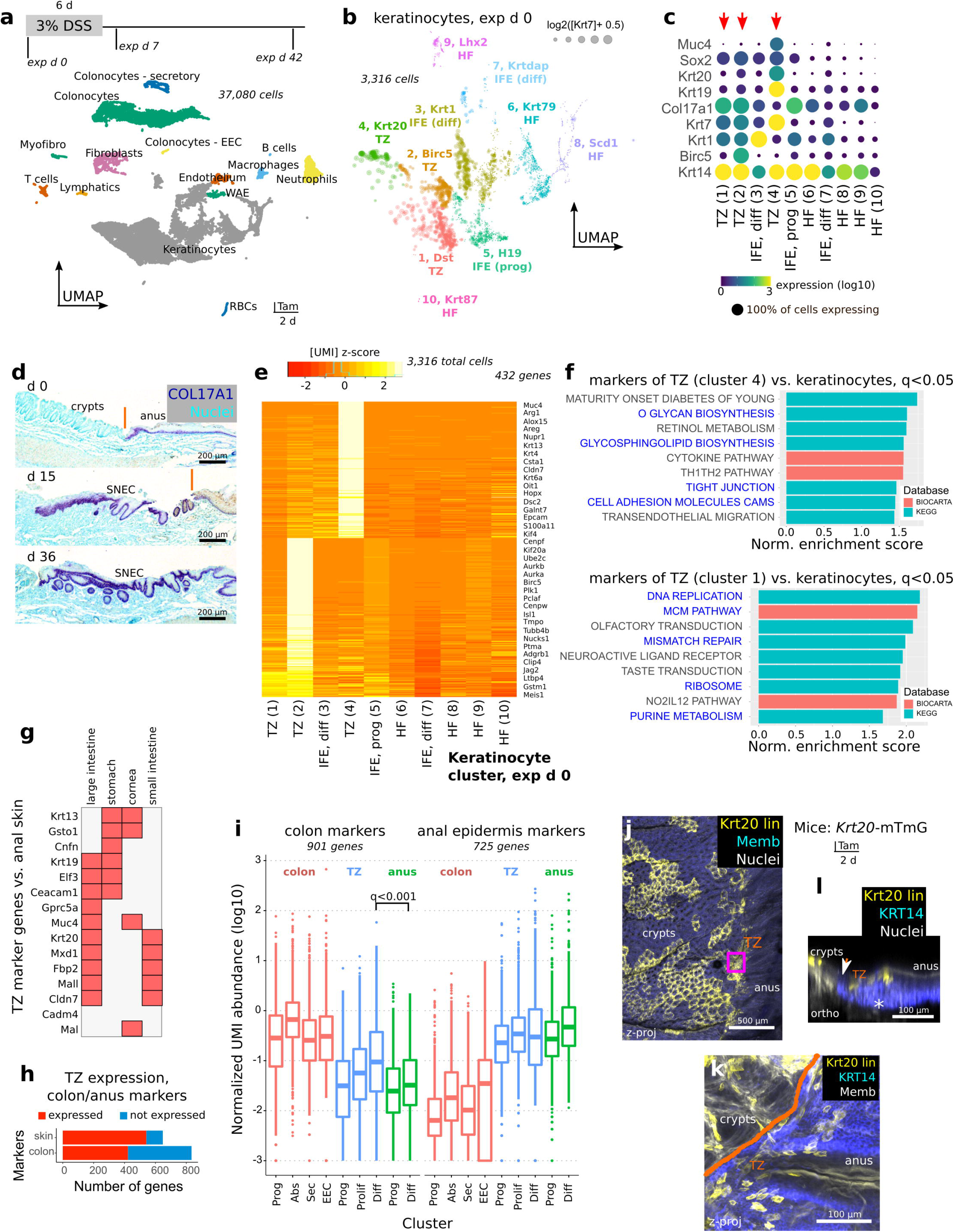
Anal TZ is a molecularly distinct squamous tissue with colonic expression. a) Single-cell RNA sequencing (scRNA-Seq) of >35,000 cells from 3 different timepoints (n=5 pooled mice/timepoint) during DSS-induced injury and repair. Discrete clusters of epithelial and stromal cells from the anorectal junction can be identified in the UMAP visualization. b-c) Clusters (algorithm: Louvain) of Krt7^hi^ cells (#1, #2, #4) representing putative TZ cells (b) and expressing specific markers among keratinocytes (c) are found at exp d 0 (no injury). d) COL17A1 staining shows that TZ, anal epidermis, and SNEC have a basal layer that is marked by expression of this collagen isoform. e) Heatmap of aggregated gene expression compared between clusters shows specific genes to clusters #1, #2, and #4 representing unique TZ-related transcripts. f) GSEA of markers of cluster #4 (differentiated TZ) and #1 (basal TZ) identifies junctional and mucus-producing pathways (#4) and mitotic pathways (#1) (blue text) enriched in the TZ. Normalized enrichment score and adjusted p values (q values) were obtained from GSEA software. g) Specific markers of the TZ cross-compared to other tissues using BioGPS. h,i) Summary (h) of colonic or anal epidermal (skin) marker expression in the TZ. By definition, skin and colonic markers were mutually exclusive. The boxplot (i) shows the distribution of expression of colonic or anal epithelial markers in different colonic or anal epithelial cell types. Each dot represents a gene. Expression is quantified as the mean UMI abundance (including the 0-count cells) over the cells within a cluster. Colon and anus markers were defined by identifying genes that had at last 4-fold expression difference between colonic and anal epithelium (the TZ was excluded for gene identification). The “differentiated” TZ cells exhibit significantly enriched overall expression of colonic epithelial markers compared to differentiated cells of the anal IFE. Significance was evaluated using a paired t-test with Holm-Bonferroni correction. In (h), a marker was deemed expressed in the TZ if its mean log TZ abundance was >50% of the abundance in the reference tissue. Abbreviations: Prog = progenitors, Abs = absorptive cells, Sec = secretory cells, EEC = enteroendocrine cells, Diff = differentiated cells. Box plot: center line, median; hinges, 25-75^th^ percentiles; whiskers, 1.5x interquartile range; points, outliers. j-l) Krt20 expression in the uninjured anal TZ was confirmed using Krt20-mTmG mice (n=3 mice). Tamoxifen administration induces lineage labeling of Krt20+ cells in colon and in the TZ (j). TZ-labeled cells distal to the dentate line (orange) are KRT14+ in the zoomed image (k) and are restricted to the suprabasal layer of cells (l). See also Figures S8, S9.

We next subclustered the Krt14^hi^ keratinocytes at exp d 0, prior to injury (Figure S8d,e). We identified 3 small clusters (#1, #2, #4) of squamous (Krt14+) Krt7^hi^ cells that lacked squamous terminal differentiation markers such as Krt1 (Figure 4b). Clusters #1 and #2 both expressed high levels of the squamous basal cell marker Col17a1, but cluster #2 also expressed mitotic markers such as Birc5 (survivin) (Figure 4c,d). We conclude that these clusters contain proliferative and nonproliferative basal cells of the TZ. In contrast, cluster #4 was negative for Col17a1 and Birc5, suggesting it represents suprabasal TZ cells. These suprabasal TZ cells were a unique differentiated cell population, with a distinct gene expression profile, compared to keratinocytes of the anal epidermis and the hair follicle (Figure 4e). Pathway analysis of cluster #4 marker genes demonstrated enrichment of tight junction and mucus-processing pathways. In contrast, the basal layer of TZ cells (cluster #1) was elevated for mitotic transcripts (Figure 4f). In the TZ differentiated cluster #4, we noted specific expression of Muc4 and Krt20, markers of colonic epithelium. By comparing markers of cluster #4 to external databases of tissue-specific expression, we found a number of colonic genes that were upregulated in the TZ (Figure 4g). To formally evaluate the similarity between the TZ and colonic epithelium, we identified >1,600 mutually exclusive markers of colonic or anal epithelium from our dataset, and then assessed their overall expression level in the TZ. The TZ expressed markers of both colonic and anal epidermal epithelium (Fig. 4h). The expression of colonic-associated transcripts was significantly higher in TZ than in the anal epidermis (Fig. 4i). We obtained similar results using a graph abstraction technique to evaluate similarity between clusters (Figure S9). We next histologically validated the expression of a colonic marker in the squamous TZ. Using short-term labeling (2 d trace) in Krt20-mTmG mice (McMahon et al., 2008), we identified mosaically labeled Krt20+ cells in colonic crypts and in the non-proliferative suprabasal layer of the anal TZ, but not in the anal skin (Figure 4j-l). Thus, the uninjured TZ represents a specialized hybrid zone of squamous tissue that expresses markers of colonic epithelial differentiation.

### TZ-derived cells adopt a progenitor state during SNEC formation

In colonic epithelium, wound healing is associated with the adoption of an interferon-induced fetal-like progenitor state (Nusse et al., 2018; Yui et al., 2018). Using the scRNA-Seq data, we tested whether a similar process underpinned squamous cell-mediated colonic re-epithelialization. We specifically sub-clustered Krt7^hi^ (TZ-derived) cells across all timepoints (Figure 5a). We found that Krt7^hi^ cells obtained from exp d 7 predominantly segregated as a distinct cluster, #3. We refer to this cluster as wound-associated squamous cells (WASCs) (Figure 5b). This cluster was like cluster #1 in its expression of the basal marker Col17a1 (Figure 5c). However, WASCs were enriched for the wound-associated keratin Krt17, the interferon target Ly6a (SCA-1), and the urea-processing enzyme Ass1, among other genes (Figure 5c). SCA-1 (Ly6a) protein and Ass1 mRNA expression, markers of the WASCs of cluster #3, were histologically confirmed in the leading edge of basal SNEC cells during wound healing (Figure 5d,e). Direct gene set enrichment analysis (GSEA) of the global expression profile of WASCs versus other Krt7^hi^ cells showed enriched translational and ribosomal pathways and similarity to stem cell states (Figure 5f). Thus, WASCs exhibit progenitor-like signaling.

**Figure 5:**
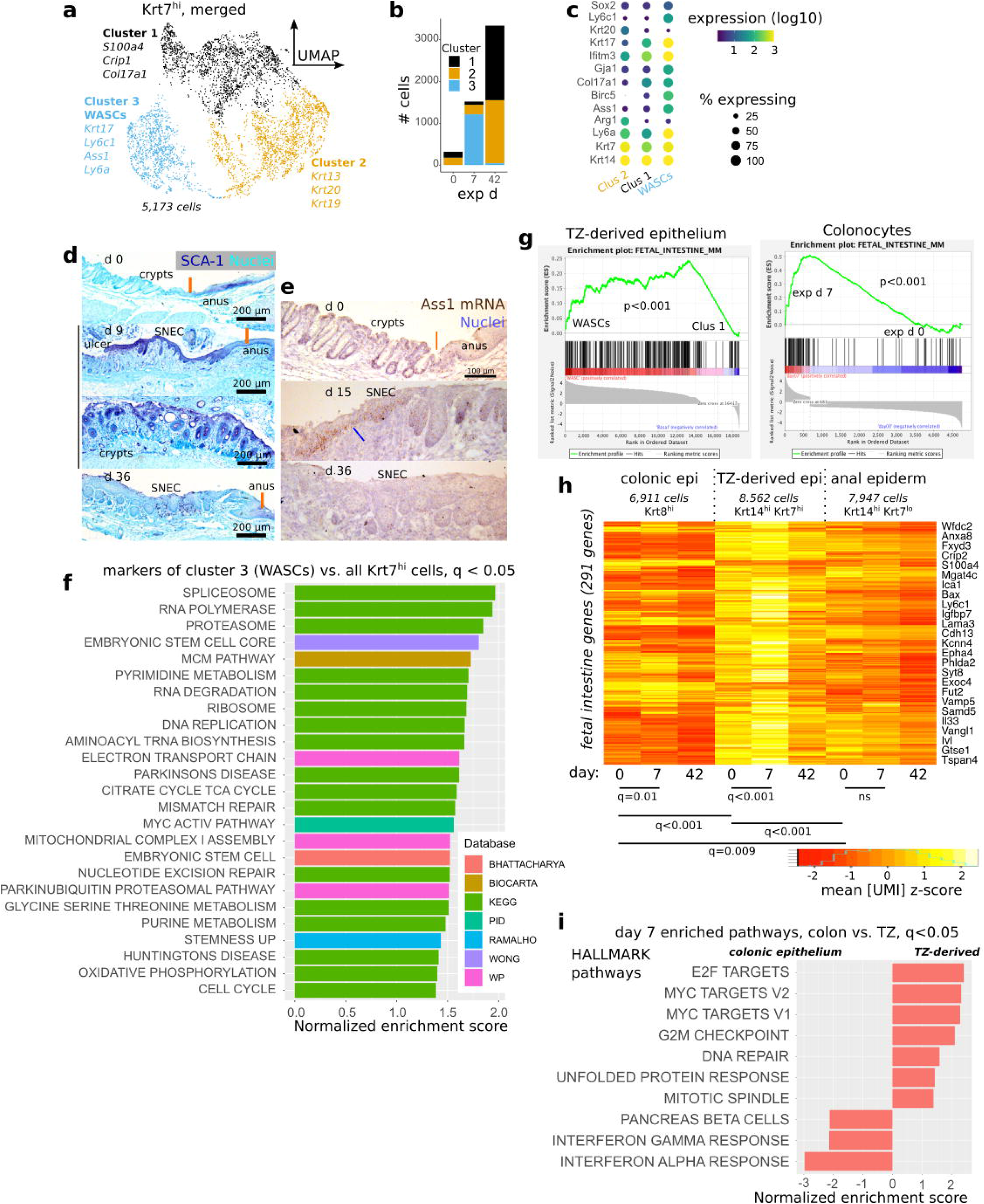
TZ-derived cells adopt a progenitor state during SNEC formation. a) Clustering of Krt7^hi^ cells combined from all 3 timepoints analyzed via scRNA-Seq; UMAP visualization reveals a distinct cluster (#3) containing wound associated squamous cells (WASCs) from exp d 7. b) Plot of cell count per cluster shows that cluster #3 predominantly mapped to Krt7^hi^ cells at exp d 7. c) Enriched genes in WASCs include Ly6a, Krt17, and Ass1. d) Leading edge cells in SNEC at exp 9 exhibit high expression of SCA-1 (Ly6a) protein, similar to colonic crypt cells involved in wound repair. SCA-1 expression is reduced prior to injury and after repair. e) Leading edge cells (blue line) in SNEC selectively express Ass1 mRNA during injury. f) Significant pathways from gene set enrichment analysis of WASC marker genes demonstrates ribosomal, translational, and stem cell pathways. g) Enrichment plots from TZ-derived epithelium and colonocytes demonstrates that genes related to fetal intestinal epithelium are upregulated in WASCs and reprogrammed colonocytes. h) Heatmap of aggregate fetal intestinal gene expression in different types of epithelial cells at exp d 0, 7, or 42. Significance was evaluated using paired t-tests with Holm-Bonferroni correction. The adjusted p value is reported as a “q value.” i) Significant HALLMARK pathways from GSEA comparing the effect of DSS-induced injury on colonic epithelium vs. TZ-derived cells at exp d 7. The inflammatory response is more pronounced in colonic epithelium.

The Ly6a transcript upregulated in WASCs is a marker of intestinal fetal epithelium (Nusse et al., 2018; Yui et al., 2018). We therefore directly assessed whether the fetal intestinal profile was upregulated in WASCs. Using GSEA, we demonstrated significant upregulation of this pathway in both WASCs and colonocytes at exp d 7 (Figure 5g). Moreover, comparison of the overall abundances of fetal genes across colonic, TZ-derived, and anal epidermal tissues showed that injury-induced fetal enrichment was specific to colon and TZ-derived cells, but not the anal epidermis (Figure 5h). The TZ-derived cells and to a lesser extent the anal epidermis exhibited a higher baseline (non-injury) expression level of intestinal fetal genes. This suggests that, at baseline, the intestinal fetal gene set contains genes associated with squamation. Despite similar responses in fetal genes, TZ-derived epithelium functionally diverged from colonic epithelium at exp d 7, likely due to differences in the response to DSS. The TZ-derived tissue was transcriptionally enriched for proliferative pathways, whereas the colonic epithelium expressed hallmarks of cytokine-induced signaling (Figure 5i). This is consistent with the overall difference in BrdU incorporation between the two epithelia (cf. Figure 3b) in early healing. In total, these results show that anal TZ cells are transcriptionally reprogrammed to a progenitor status similar to the fetal intestine during SNEC formation.

### SNEC upregulates colonic expression patterns

Prior to injury, anal TZ cells reside at the anorectal squamocolumnar junction and have mixed squamous and intestinal gene expression patterns. However, the injury-induced derivative tissue, SNEC, exists within the colon. Because epithelial stem cells can both rely on and leverage local microenvironmental cues, we hypothesized that SNEC cells would exhibit markers of adaptation to the colonic milieu. We therefore tested whether SNEC cells further upregulate colonic expression modules. Focusing on the suprabasal cluster (#2) shown in Figure 5a, we found evidence of altered transcriptional profile in SNEC (exp d 42) versus the uninjured TZ (exp d 0) (Figure 6a). We performed pathway analysis of differentially expressed genes identified through regression analysis of SNEC vs. TZ. We found that SNEC upregulated a tight junction module and other colonic transcripts, while downregulating epidermal transcripts (Figure 6b). The expression of individual claudins and colon-specific keratins was significantly elevated in SNEC vs. TZ (Figure 6c). Thus, SNEC elaborates the hybrid colonic-epidermal profile of the TZ and becomes more colonic. At the global level, SNEC retained some similarity to skin tissue but had significantly elevated expression of colonic genes (Figure 6d). As a hybrid epithelium, SNEC exhibited a distinct differentiated cell profile. Suprabasal cells of SNEC immunoreacted with antibodies targeted against the colonic marker MUC4, a mucus-associated protein, the colonic tight junction protein claudin-1 (CLDN1), and GSTO1, a specific glutathione transferase found in SNEC (Figure 6e, Figure S10). To determine whether the specific suprabasal differentiation of SNEC was supported by a uniquely marked basal cell layer, we analyzed and re-clustered all SNEC (Krt7^hi^) cells at exp d 42 (Figure 6f). We identified a suprabasal Krt20+ cluster and a basal (Col17a1^hi^) proliferative (Birc5^hi^) cluster (Figure 6g). These basal cells were unique to the TZ and SNEC through their expression of Krt17 (Figure 6h,i). Collectively, these results demonstrate that SNEC is a distinct tissue type with shared features of colonic and epidermal expression.

**Figure 6:**
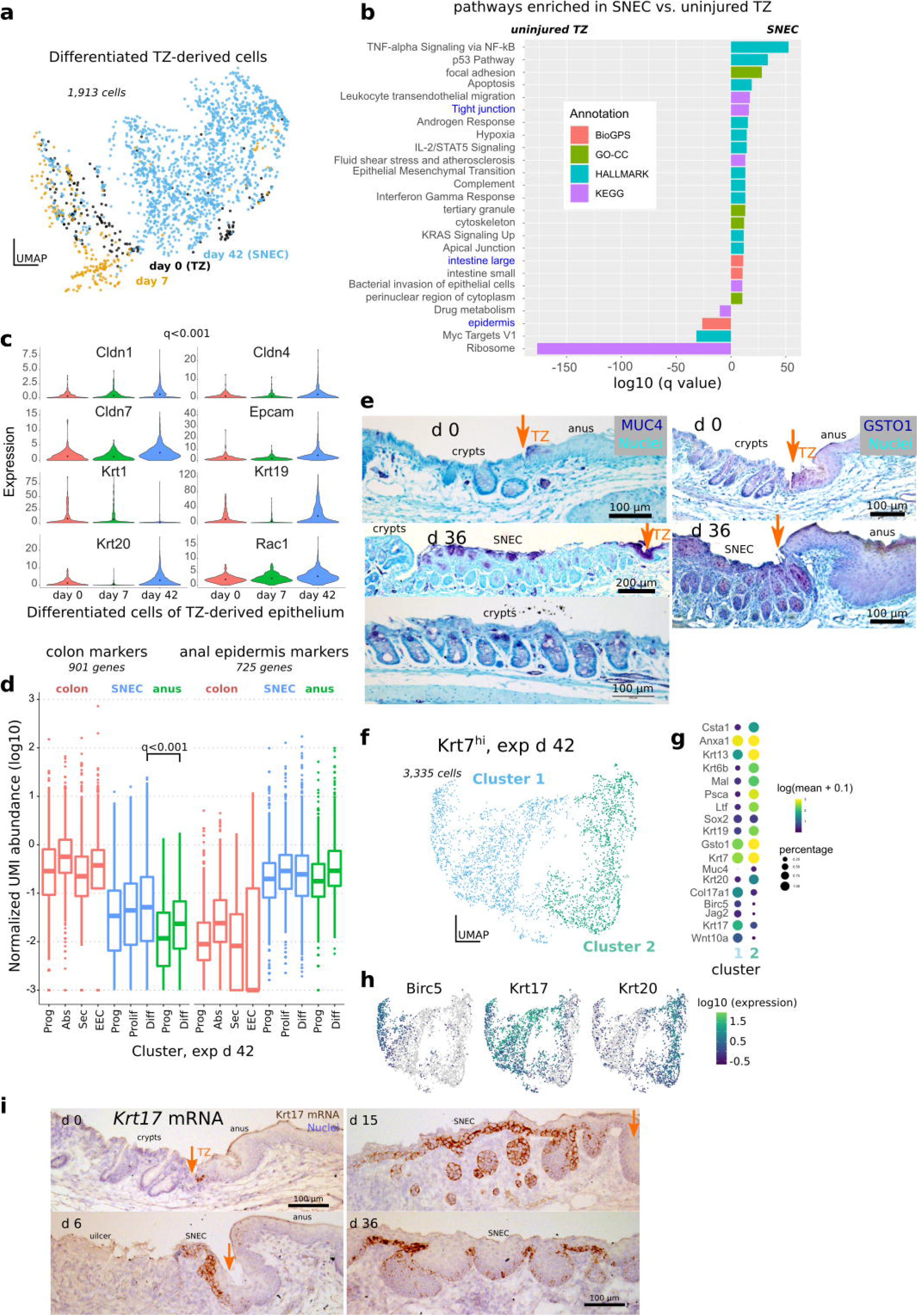
SNEC upregulates colonic expression patterns. a) UMAP of differentiated (suprabasal, Col17a1^lo^) Krt7^hi^ cells coded by experimental day. Datapoints representing SNEC cells at exp d 42 appear to be displaced from datapoints of TZ-derived cells at exp d 0 and exp d 6. b) Comparison of enriched genes at exp d 42 include those associated with tight junction pathways and colonic tissue. Annotations are obtained using enrichr. c) Violin plots of significant increases in junctional and colonic genes at exp d 42; significance was evaluated by the fit of a sloped linear regressor to the expression values. d) Boxplot similar to that shown in Fig. 4h comparing expression of colonic and anal marker expression at exp d 42. Like uninjured TZ, SNEC has elevated expression of colonic markers compared to anal epidermis. e) Immunostaining for MUC4 shows presence of colon-like “differentiated” cells in SNEC in the suprabasal layer at exp d 36. MUC4+ cells were also found in the uninjured TZ. GSTO1-targeted antibody labels TZ and SNEC suprabasal cells, but not crypt cells. Orange arrows mark the dentate line. f) UMAP of TZ-derived cells at exp d 42 shows 2 distinct clusters. g) The dotted plot indicates a proliferative (Birc5^hi^) basal (Col17a1^hi^) cluster (cluster 1) and a nonproliferative suprabasal cluster enriched for Krt19, Krt20, and Muc4. h) Expression of proliferative markers (Birc5) and specific keratins (Krt17, Krt20) in different cells at exp d 42. i) In situ hybridization reveals that Krt17 mRNA expression specifically marks basal cells of the uninjured TZ and of SNEC. See also Figure S10.

### SNEC harbors stem cells that mediate tissue homeostasis in a crypt-like pattern

In addition to upregulation of colonic transcripts, SNEC is composed of crypt-like rete peg structures. To assess further evidence of the adoption of colonic-like features, we tested whether rete pegs functioned similarly as crypt-like units of regeneration. To study the patterns of cell turnover and regeneration in SNEC, we turned to long-term clonal lineage tracing using Sox2-Confetti mice. Over a 28-wk time course, we found that a growing proportion of rete pegs became fully monoclonal, comprising cells marked by a single Confetti color (Figure 7a). Thus, once established, rete pegs function as a closed renewing glandular-like compartment, similar to the epithelial organization of the uninjured colonic crypt. More generally, 3 types of clones could be resolved in the tracing data: (1) clones occupying the lateral boundary of the rete peg (termed “casing” clones); (2) clones extending from the abluminal “tip” (i.e., basal vertex) into the inner, non-cycling “core”; and (3) monoclonal rete pegs (Figure 7b,c). Similar results were obtained from the analysis of patches of nGFP expression, which recombines with ~10% of the efficiency of the other Confetti colors (Snippert et al., 2010), allowing us to conclude that spatial variability was not the result of clone merger events. The spatial segregation and relative persistence of these clonal types suggested sub-compartmentalization of the epithelium, with a closed lateral boundary domain and an inner core maintained by proliferative cells at the basal tip. We reasoned that an additional transfer of cells from the tip to the casing region would account for the clonal fixation of rete pegs in the long term (Figure 7d). These results suggest that SNEC is renewed in the long-term by the lineages of stem cells located at the rete peg base.

**Figure 7:**
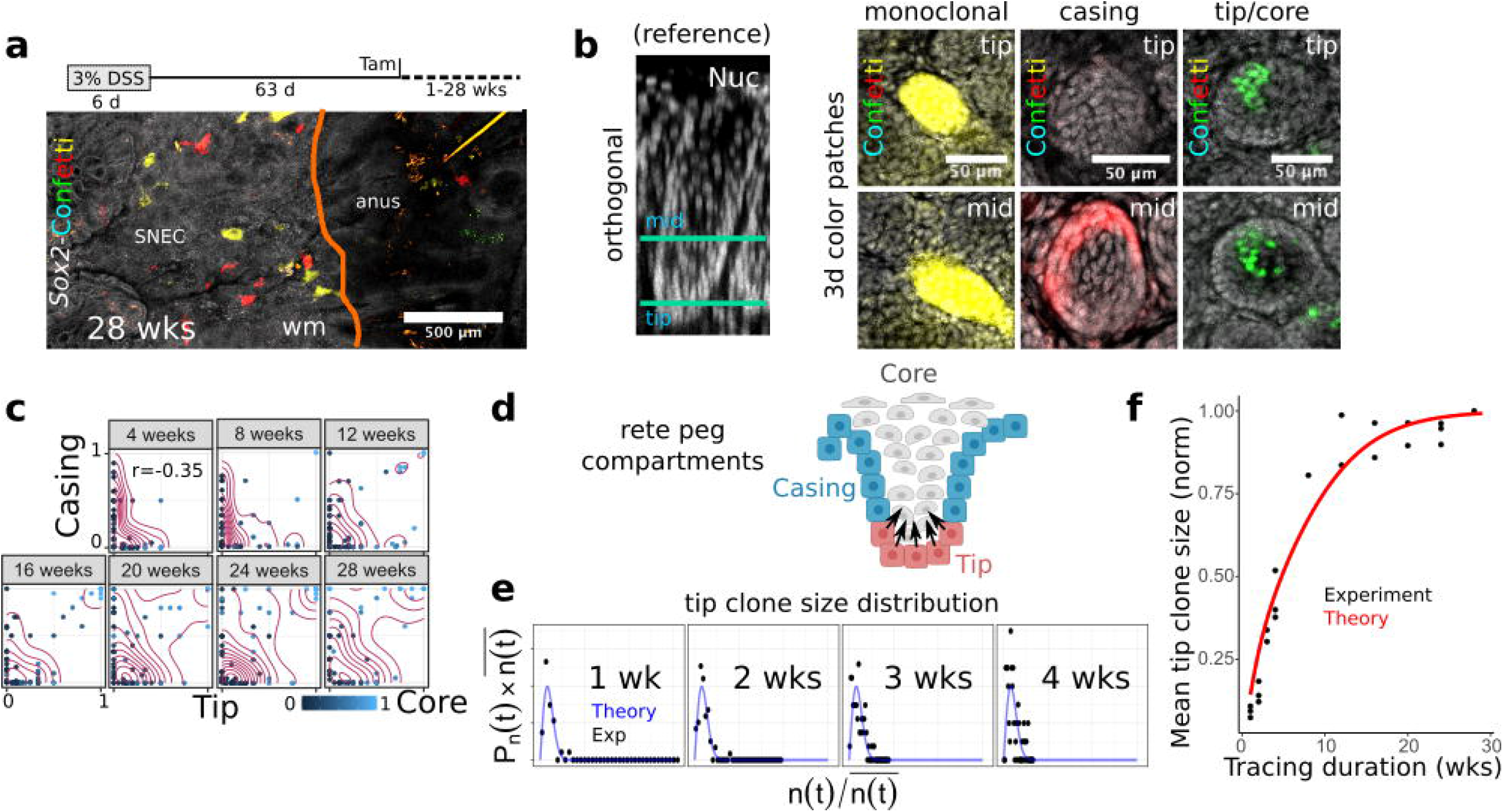
SNEC harbors stem cells that mediate tissue homeostasis in a crypt-like pattern. a) Long-term clonal tracing (28 wks) in Sox2-Confetti mice demonstrates clonal fixation (single color) in rete pegs (n=3 mice/timepoint). b) Example images of clonal forms (color patches) identified during 1-28 wks of lineage tracing. A reference rotated xy-image of a rete peg is shown to orient the individual xz-planes demonstrating the fate-mapping. “Tip” and “mid” describe lowest and middle height in the rete peg, respectively. c) Quantification of clone stratification between tip, casing, and core domains demonstrates no correlation in clone size between tip and casing compartments, suggesting that progenitor cells in these compartments function independently. Tip and core clone sizes are correlated (coeff = 0.74). d) Diagram of predicted progenitor cell populations in the rete peg. e) Superimposing the 1d scaling form to a plot of the mean-normalized tip clone sizes (n(t)) vs. their frequency (P_n_(t)), obtained from tracing data from 1-4 wks, shows a good fit, demonstrating neutral drift dynamics in tip progenitor population. f) Fit of neutral drift model to the empirical normalized mean tip clone size over time (λ/N^2^=0.02 /wk), consistent with theory of replacement dynamics of an equipotent progenitor cell population. See also Figures S11, S12.

Maintenance of the intestinal epithelium relies upon the constant loss and replacement of equipotent crypt base columnar cells leading to neutral drift clone dynamics, culminating in clonal loss or fixation of the crypt (Kozar et al., 2013; Lopez-Garcia et al., 2010). We next assessed whether neutral drift dynamics also characterize turnover of tip progenitor cells. In tracing initiated with low doses of tamoxifen, we found that the clone size distributions obtained from the independently functioning tip and casing compartments conformed to a statistical “scaling” behavior over the short-term (4 and 7 d) (Figure S11). Over 4 wks of tracing, the tip clone size distribution resembled a statistically time-invariant function whose spread was dictated by the mean, a hallmark of scaling behavior (Figure 7e). Moreover, over the full duration (28 wks) of tracing, we found that the empirical mean surviving tip clone size was well fit by a neutral drift model (Figure 7f), with a key parameter (*λ*/*N^2^* = 0.02 /wk) describing scaling dynamics similar to previously reported values (Lopez-Garcia et al., 2010) in colon and small intestine (see Methods, Figure S12). Thus, we reasoned that the rete pegs are sustained by an equipotent progenitor cell population at the abluminal tip. This progenitor cell population operates from an analogous invaginated location and with similar dynamic features (Lopez-Garcia et al., 2010) as colonic epithelial stem cells.

## Discussion

Colonic epithelial damage represents an immediate crisis requiring rapid repair. Columnar epithelial cells can undergo dedifferentiation to promote healing (Murata et al., 2020; Radyk et al., 2018; Tetteh et al., 2016). Here, using high-resolution imaging, single-cell transcriptomics, and biophysical modeling, we show that severe colitis is also associated with a previously undescribed plasticity response: the formation of a permanent squamous epithelium, SNEC. This represents a regenerative process that engages a heterologous cell type. Unlike the anal skin, the injury-associated squamous epithelium remodels to upregulate key features of colonic columnar epithelium: morphological adoption of crypt-like units in the form of rete pegs, expression of differentiated colonocyte markers including certain mucins, keratins, and junctional proteins, and long-term regeneration through a stem cell population with intestine-like localization and dynamics. These are key features that make SNEC suitable for colonic wound healing. SNEC represents a colonic extension of an anal transition zone that exhibits a hybrid of colon-epidermis features in homeostasis. This squamous epithelium expands into the colon because it is partially resistant to the severe injury and inflammation that can induce proliferative arrest in colonocytes. Consistent with this paradigm, repetitive or chronic injury was associated with larger areas of SNEC. We do not know if there is a proximal limit to the contiguous expansion of SNEC. Given its overall cellular plasticity and its ability to thrive in the heterogeneous microenvironment of the anal TZ, which supports both colonic and epidermal tissues, SNEC could theoretically move into the proximal (ascending) colon.

SNEC bears resemblance to a squamous metaplasia described in case reports of human patients with ulcerative colitis or distal colonic injury (Adamsen et al., 1988; Bujanda et al., 2001; Cheng et al., 2007; Mahesha et al., 2006; Nishi et al., 2004; Pantanowitz, 2009); however, accurate estimates of prevalence in humans are challenged by lack of sampling at the anorectal junction during endoscopic screening, and limited surgical samples of completion proctectomy with removal of the anal canal. The underlying processes giving rise to SNEC may be similar to those of early metaplasia. For example, both SNEC and Barrett’s esophageal metaplasia exhibit elevated expression of Krt7 (Cabibi et al., 2009) and share an injurious ontogeny. Whether SNEC can be further directed to an intestinal lineage like Barrett’s metaplasia remains unknown. Intestinalization of SNEC could support its longer-term role in the restoration of colonic function. However, retention of a squamous profile may be integral to providing resistance to colonic injury. In addition, future studies are needed to assess the relationship between SNEC and cancer, a sequela of metaplasia. This could inform on the physiological trade-offs between the rapid re-epithelialization of the colon by an injury-resistant squamous tissue and a long-term potential for carcinogenesis.

SNEC is derived from a specialized population of squamous cells located within the small transitional zone of the anorectal junction. The width of the TZ is <10 cells-thick by the marker of Krt7 mRNA expression; KRT7 antibody staining and lineage tracing in Krt7::CreER transgenic mice show a marginally wider domain. This discrepancy could be explained technically by differing sensitivities of the detection assays, or explained biologically through the lateral spread of the descendants of Krt7^hi^ TZ cells. Though few in number, anal TZ cells represent a distinct tissue type that co-expresses high levels of Krt7, Sox2, and Krt14. A transition zone has been identified at the gastroesophageal squamocolumnar junction; these cells can be directed to express intestinal markers in vitro (Jiang et al., 2017). Thus, transitional epithelia in the body may broadly represent important areas of plasticity. The colonic expression pattern of SNEC is an extension of a pre-existing colon-like expression profile in the anal squamous TZ. An intriguing observation of the anal TZ is the baseline expression of Krt17 (Figure 6i), considered a wound-associated keratin (Zhang et al., 2019), and the overall similarity between intestinal wound healing and squamation (Figure 5h). Given the rapid migration and proliferation of TZ cells during distal colonic ulceration, the TZ may represent a reserve set of cells that are primed to respond to colonic injury. Because of the dichotomous microenvironment at tissue junctions, any epithelium at these junctions must have some inherent adaptability. It is therefore evolutionarily beneficial that a wound-adapted cell population would reside at these specific sites to quickly respond during tissue insults.

Squamous cell phenotypes appear to be important for optimal outcomes after colonic injury. Colonic epithelial cells involved in wound restitution adopt a flattened morphology that is reminiscent of squamation (Miyoshi et al., 2012). This study complementarily demonstrates the importance of anal-derived squamous Sox2+ cells. Mice with increased apoptosis within Sox2+ cells could not properly recover from acute colitis. Given the role of Sox2+ cells in mediating the re-epithelialization through the formation of SNEC, a simple explanation for this observation is the impaired restitution by anal-derived squamous cells. Some squamous cells of SNEC persisted even with apoptotic targeting; we therefore cannot rule out that squamous cells have a secondary modulatory impact on the extent of injury or survival of colonic epithelium. For example, TZ cells express amphiregulin (Areg) (cf. Figure 4e), an EGFR ligand that can have both pro-repair and anti-inflammatory functions on the wound microenvironment (Zaiss et al., 2015).

A deeper understanding of the diverse cell-types and mechanisms involved in colonic epithelial repair is of therapeutic relevance to chronic inflammatory diseases such as IBD. The identification of a unique population of junctional skin-like cells capable of rapid colonic restitution supports the potential of heterologous squamous cell phenotypes to serve as targets for mucosal healing. This illuminates new strategies for the design of heretofore elusive therapies that directly activate regenerative processes in the colon.

## Acknowledgments

This work was supported by the U.S. National Institutes of Health (R01DK108648 and R01DK056008 awarded to D.B.P.), the Crohn’s and Colitis Foundation (Career Development Award 694110 to C.Y.L.), and the California Institute for Regenerative Medicine (postdoctoral fellowship to C.Y.L.). B.D.S acknowledges funding from the Wellcome Trust (098357/Z/12/Z) and the Royal Society through an EP Abraham Research Professorship (RP/R1/180165).

## Author Contributions

C.Y.L. and D.B.P. conceived the project and supervised all project-related activities. C.Y.L., N.G., M.L.G., and P.E.D. collected primary data. M.K.W. contributed analysis of histological data. C.Y.L. and B.D.S. analyzed lineage tracing data. C.Y.L. and D.B.P. interpreted all experimental data. C.Y.L. wrote the manuscript. B.D.S. and D.B.P. edited the manuscript. All authors approved the final version of the manuscript.

## Methods

### Resource availability

Key resources used in the study are listed in the Table.

#### Lead contact

Further information and requests for resources and reagents should be directed to and will be fulfilled by the lead contract, D. Brent Polk.

#### Materials availability

This study did not generate new unique reagents.

#### Data and code availability

Bulk RNA sequencing data shown in Figure S3 are available on the Gene Expression Omnibus (GEO), with accession number GSE168053: https://www.ncbi.nlm.nih.gov/geo/query/acc.cgi?acc=GSE168053. Single-cell RNA sequencing data shown in Figs. 4–6 are available on GEO, with accession number GSE168033: https://www.ncbi.nlm.nih.gov/geo/query/acc.cgi?acc=GSE168033. Analysis scripts for singlecell sequencing are available on GitHub: https://github.com/stalepig/2021-SNEC-single-cell. Imaging processing routines for images obtained via the Deep Mucosal Imaging (DMI) pipeline are available on GitHub: github.com/stalepig/deep-mucosal-imaging. All other datasets generated and analyzed during the current study are available from the lead contact on reasonable request.

### Experimental model and subject details

#### Mice

All animals were maintained ethically and humanely at Children’s Hospital Los Angeles (CHLA). The Institutional Animal Care and Use Committee of CHLA approved the study under internal protocol #288. Sox2::CreER (stock 017593), Rosa26::Confetti (stock 013731), Rosa26::LSL-DTA (stock 009669), Rosa26::mTmG (stock 007676), and C57Bl6/J (stock 000664) mice were purchased from Jackson Laboratory. Vil1::Cre mice(el Marjou et al., 2004) were a kind gift from Sylvie Robine and Daniel Louvard (Institut Curie). Krt7::CreER mice (Jiang et al., 2017) were generously provided by Jianwen Que (Columbia). Krt20::CreER mice (McMahon et al., 2008) were donated by Andrew McMahon (University of Southern California).

Mice were housed in the Animal Care Facility at CHLA. The mice were provided with standard bedding and airflow. Irradiated food pellets and drinking water supplied from a valve were provided for consumption ad libitum. The housing room was kept on a strict 12 h light/dark cycle and maintained at 20 °C. Littermate mice were separated by sex at 3 wks-old and co-housed to a maximum of 5 mice per cage. Lineage tracing and immunolabeling experiments were performed on both male and female transgenic mice. For sequencing studies, cohorts of cohoused 6-wk-old male wildtype mice (C57Bl/6J) were ordered from Jackson Laboratory and allowed to habituate to the animal room for 3 wks prior to experimentation.

Retrospective analyses of Il10-/-mice were performed from archived slides of a previous study (Dube et al., 2012).

### Method details

#### Dissection and fixation

For experimental studies, mice were anesthetized with isoflurane and euthanized via cervical dislocation. The colon and adjacent anal ring were removed, opened longitudinally, and washed of fecal material. Tissue was immediately flattened against blotting paper and fixed overnight in methanol-free 4% cold paraformaldehyde/phosphate-buffered saline (PBS). The next day, the mucosal layer was finely freed under a stereomicroscope from serosal/muscular layers and additionally processed.

#### Deep Mucosal Imaging (DMI)

The pipeline for mounting the specimen, imaging, and in silico reconstruction of images was as previously published (Liu et al., 2015; Liu and Polk, 2020). However, the clearing reagents used are different. Briefly, fixed tissues were washed with PBS, incubated with 20% sucrose/PBS and frozen. Tissues were thawed and incubated with CUBIC-1 (Susaki et al., 2014) clearing agent supplemented with 1:1,000 dilution of a 4% w/v methyl green stock (Prieto et al., 2015) that was purified of contaminating crystal violet through multiple phase separations in chloroform. Tissues were cleared for 1-7 d prior to imaging. For imaging, tissues were embedded between two #1 glass coverslips and mounted with the used clearing solution. Tiled images were captured on a Zeiss LSM 700 confocal microscope with either 10X/0.45NA or 20X/0.8 air objectives. The distal 1-3 cm of mouse colonic mucosa and adjacent anal tissue were captured in overnight scans.

#### Whole-mount antibody staining

For procedures not requiring the preservation of fluorescent proteins, the iDISCO procedure (Renier et al., 2014) was modified for mouse colon. Fixed tissue was washed with PBS and permeabilized in an ascending (20-100%) methanol/PBS series, and then incubated overnight in 5% hydrogen peroxide/methanol at 4°C. Tissue was rehydrated in a descending methanol/PBS series, washed successively with PBS and 0.3% Triton X-100 in PBS (PBTx), and incubated for 18 h at 37°C in 20% DMSO/0.3M glycine in PBTx and for 24 h at 37°C in 10% DMSO/5% normal horse serum prepared in PBTx. Primary antibody was diluted in PBTx supplemented with 10 μg/ml heparin, 3% normal horse serum, and 5% DMSO. Samples were stained with primary antibody at 37°C for 4 d. Washes were performed with PBTx with 10 μg/ml heparin. Secondary antibodies conjugated with Alexa dyes (Thermo Fisher Scientific) were diluted in PBTx with 10 μg/ml heparin, 3% normal horse serum, and a 1:10,000 dilution of 4% w/v methyl green stock. Secondary staining was performed for 3 d at 37°C. Stained samples were washed and chemically cleared in CUBIC-2 for 1-7 d and mounted, imaged, and reconstructed using the DMI pipeline.

Antibodies used in the iDISCO protocol were rabbit anti-cleaved caspase-3 (Cell Signaling Technology 9661, 1:100) and rat anti-KI-67 (eBioscience 41-5698, clone SolA15, 1:100).

For procedures requiring the preservation of fluorescent proteins, we first cleared the tissue using CUBIC-1 for 3 d at room temperature. The tissue was washed with PBS and blocked at room temperature overnight in 0.3% PBTx supplemented to 5% with horse serum. Primary antibodies were diluted in blocking solution and incubated with the tissue for 3 d at room temperature. Tissue was washed with PBS. Secondary antibodies were diluted in 0.3% PBTx and incubated with tissue for 3 d at room temperature. After a final series of washes with PBS, tissue was cleared with CUBIC-2 for 1-4 d, mounted, and directly imaged in the clearing solution. Primary antibodies were directed against KRT7 (Abcam ab181598, 1:200) and KRT14 (Abcam ab206100, 1:500).

#### In situ hybridization

Chromogenic in situ hybridization was performed using the RNAScope 2.5 HD Assay – Brown (ACDBio/Biotechne) according to the manufacturer’s instructions. Paraffin-embedded sections were hybridized with probes targeted to murine *Krt17* or *Ass1*. Signal was revealed with DAB (Vector Labs), and the tissue was counterstained with hematoxylin and blued with ammonium hydroxide.

Fluorescent in situ hybridization was performed on paraffin-embedded sections with hybridization chain reaction (HCR) technology (Molecular Instruments) according to the manufacturer’s recommendations. Probes were directed against *Krt7*. After staining, samples were mounted in Fluoro-Gel (Electron Microscopy Sciences) prior to confocal imaging.

#### Histochemistry

Immunohistochemical experiments on paraformaldehyde-fixed, paraffin-embedded or frozen sections was performed according to standard procedures. For paraffin-embedded sections, a citric acid/heat antigen retrieval step was performed prior to antibody incubation. Endogenous peroxidases were quenched with a 30 min incubation in 3% hydrogen peroxide in PBS. Primary antibodies used were rat anti-CDH1 (R&D Systems FAB7481F, paraffin, 1:300), rabbit anti-P63 (Proteintech 12143-1-AP, paraffin, 1:200), rabbit anti-SOX2 (Abcam ab97959, paraffin, 1:1500), rabbit anti-CLDN1 (Thermo Fisher Scientific 71-7800, paraffin, 1:1000), rabbit anti-LOR (Biolegend 905101, 1:5000), mouse anti-MUC4 (Thermo Fisher Scientific 35-4900, 1:2000), rabbit anti-KRT17 (Sigma-Aldrich AV41733, 1:3000), rabbit anti-COL17A1 (Abcam ab184996, 1:250), rabbit anti-GSTO1 (Novus NBP1-33763, 1:100), rabbit anti-SCA-1 (Abcam ab109211, 1:2000), rabbit anti-KRT8 (Abcam ab53280, 1:1000), rabbit anti-IVL (Biolegend 924401, 1:10000), rabbit anti-TGM1 (Proteintech 12912-3-AP, 1:500), rabbit anti-KRT1 (Biolegend 905602, 1:1000), with HRP-linked secondaries. Chromogen development was performed using a VIP kit (Vector Labs), and slides were counterstained with methyl green.

#### Mouse endoscopy

Mice were sedated with ketamine and xylazine. Distal colonoscopy was performed with a KARL STORZ Veterinary Endoscope (Karl Storz, Tuttlingen, Germany).

#### DSS colitis

Dextran sulfate sodium (DSS)-induced colitis was administered as described previously (Liu et al., 2015). Mice were trained to drink solely from a water bottle provided to the cage for 3-7 d prior to administration of DSS (38-50 kDa). On exp d 0, the contents of the water bottle were replaced with dextran sulfate sodium supplemented to de-ionized drinking water to a final concentration of 3% w/v. After 6 d, the water bottle was removed, and the cage was returned to the housing room’s water valve supply.

#### Lineage tracing experiments

Induction of CreER proteins for lineage tracing was induced by a single injection of 2 mg tamoxifen. Sox2::CreER;Rosa26:: Confetti lineage tracing for clonal analysis was initiated with a single intraperitoneal injection of 0.5 mg tamoxifen.

#### Target cell ablation experiments

Systemic activation of DTA expression was induced with 2 mg tamoxifen administered as in lineage tracing. Local or topical activation of DTA in the distal colon/anus was induced via enema. 4-hydroxytamoxifen (4-OHT) was dissolved in a heated mixture of 65%/35% Witepsol H-15/corn oil at 0.25 mg/ml and cooled to form a suppository. A per-dose volume of 0.1 ml was intrarectally distilled through an 18-gauge blunt needle to sedated mice. Mice received 3 doses of 4-OHT; the first dose was an “induction” dose through which mice were deeply sedated by ketamine and xylazine and inverted after enema administration to maximize rectal retention. In the other “maintenance” doses, mice were sedated by isoflurane, given the enema, and allowed to recover quickly.

#### EdU/BrdU double labeling

Mice were intraperitoneally injected with 2.5 mg EdU (Cayman 20518) 24 h prior to dissection and 2.5 mg BrdU (Abcam ab142567) 2 h prior to dissection. Nucleotides were prepared as 10 mg/ml stocks in PBS.

For dual revelation of EdU and BrdU incorporation in paraffin-embedded sections, slides with tissue were de-waxed and boiled for 20 min in a pressure cooker in 10 mM sodium citrate buffer (pH 6.0) supplemented with 0.5% Tween. The Click reaction was performed for 30 min at room temperature with 2 mM CuSO_4_, 10 mM THPTA, 2 μM Cy3 azide (Click Chemistry Tools AZ119-1), and 100 mM ascorbic acid freshly prepared and diluted in Gomori buffer (pH 7.4). After the Click reaction, immunofluorescent staining with a rat anti-BrdU antibody (1:100, clone BU1/75, Abcam ab6326) was performed according to standard protocols.

For dual revelation of EdU and BrdU in full-thickness specimens, the iDISCO protocol was followed, except the EdU solution (lacking THPTA) was applied after initial permeabilization of tissue with methanol and DMSO/Triton-X. The Click reaction was performed overnight at room temperature. After washing, tissue was acid-treated with 2N HCl for 90 min at room temperature. Tissue was then stained with the anti-BrdU antibody (1:100) according to the iDISCO timings, and cleared in CUBIC-2 prior to imaging.

#### Single-cell RNA-Seq

Primary tissue was dissected from the anorectal junction with a 2-mm margin. The mucosal/surface tissue was separated from the muscle layer and digested in warmed 0.25% Trypsin-EDTA for 30 min with agitation at 160 rpm. Tissue was triturated and passed through a 100-μm mesh filter, treated with RBC lysis solution (Biolegend), washed and spun twice at 300 rcf with divalent-free HEPES-buffered saline (Liu et al., 2020) supplemented with 1% BSA, and purified for live cells using the EasySep Live/Dead magnetic selection kit (Stem Cell Technologies). Suspensions were passed through a 40-μm mesh filter to obtain single cells.

Single-cell suspensions of the combined distal colon, SNEC, transition zone, and anus were submitted representing 3 timepoints (exp d 0, 7, and 42) in mice treated with 3% DSS for 6 d (exp d 0-6). Five wildtype Bl/6 mice were pooled for each timepoint in order to obtain sufficient numbers of cells for analysis. Barcoding and library preparation was performed, allocating a separate microfluidic well per timepoint, on the Chromium (10X Genomics) platform. Targeted recovery was 10,000 cells per timepoint. Libraries were sequenced on a HiSeq 4000 machine (Illumina).

#### Bulk RNA-Seq experiments

C57Bl6/J mice were obtained from Jackson Labs and exposed to 3% DSS for 6 d. Mice were euthanized at 0, 3, 6, 9, 14, and 20 d from the beginning of DSS exposure. The distal colon and anus mucosal layers were isolated and manually separated from the underlying muscle under the guidance of a stereomicroscope. The mucosal “peel” from each mouse was divided into 4 pieces: 1) anus, 2) metaplasia (applicable from d 9 onward), 3) distal colon, or the distal-most 500-μm colonic length of colonic crypts, and 4) less-distal colon, or the piece of colon representing 1000-500-μm linear distance from the anal margin, which is overall similar to distal colon but more resistant to DSS-induced ulceration. Tissue was agitated and rendered with a manual homogenizer (Squisher, Zymo Research) in Trizol, and RNA isolation was performed according to standard procedures. Sequencing libraries were prepared from polyA-selected RNA using Illumina stranded library preparation kits and were sequenced on 3 NextSeq high-output lanes.

*

### Quantification and statistical analysis

#### Deep Mucosal Imaging (DMI)

Software routines for post-processing of images are available at github.com/stalepig/deep-mucosal-imaging Images were stitched from metadata and visualized with a custom-written interface atop of ImageJ. Refractive signal loss through the depth of the image stack was estimated from the mean intensity of cell nuclei, fit to an exponential function, and corrected in other channels as necessary. Confetti fluorophores were excited with 405 nm (mCFP), 488 nm (nGFP and YFP), and 543 nm (RFP), while methyl green was excited with a 633 nm laser.

#### Single-cell RNA-Seq

Quantification of transcript abundance per cell was obtained from cellranger 3.0 (10X Genomics). Matrix files were imported into Monocle 3 (Qiu et al., 2017). Cells with fewer than 100 UMI reads were discarded. Count matrices were merged across samples for downstream analysis. Matrices were normalized by dividing by a size factor (the sum total of counts within a cell) and then log-transformed. Dimensionality reduction was performed via principal component analysis, taking the top 50 principal components. These principal components were used for further dimensional reduction to two-dimensional space via uniform manifold approximation and projection (UMAP). This was executed using the default options in Monocle 3 for the reduce_dimension() function. Unsupervised clustering was performed from the UMAP values using the leiden algorithm, except in Figure 4b, which was clustered using the louvain algorithm. Dimensional reduction and clustering were repeated after subsetting of data.

Markers of cell clusters were identified using the top_markers() function in Monocle 3. All cells were used for marker analysis. The specificity of each marker for each cell group was computed using the Jensen-Shannon method. The significance of each marker was assessed using general linear model with a binomial family function and a likelihood ratio test between a full and simplified model of expression. The false discovery rate is reported as the adjusted p value (q value). The top 150 markers by specificity and meeting a q value < 0.001 were then uploaded into the pathway analysis aggregator enrichr (Kuleshov et al., 2016) (https://maayanlab.cloud/Enrichr/) to identify targets of organ similarity (e.g., Figure 4g) using the BioGPS database (Wu et al., 2016). These markers were also plotted as heatmaps. The top 5 markers were used for manual annotation of clusters and celltype assignment. Clusters were not merged except to facilitate display of epithelial and TZ cells as a single unit in Figure 4a and Figure S8b,d. To compare epithelial cells of different tissue types across time in aggregate (e.g., Figs. 5h, 8b), clusters were merged for Krt8+ (colonic), Krt14+ Krt7+ Sox2+ (TZ-derived), or Krt14+ Krt7-Sox2+ (anal epidermal) cells. Explicit differential gene expression analysis between samples was performed with the fit_models() function in Monocle 3; genes with an adjusted p value (q value) < 0.2 were selected for enrichr analysis as shown in Figure 6b.

Heatmaps were generated for select groups of genes by computing the average normalized counts per gene per cluster. The expression of each gene was scaled across clusters prior to display. For GSEA(Subramanian et al., 2005), the average normalized counts for all genes were exported as text files. Gene names were translated to human symbols using the Homologene database. GSEA 4.0.3 (Broad Institute) with the default options was used to determine enrichment of KEGG, Reactome, Gene Ontology, or HALLMARK pathway databases in the expression data. To assess SNEC similarity to marker genes of fetal intestinal epithelium, a dataset of gene expression from fetal mouse intestinal organoids (Mustata et al., 2013) was used as a reference.

To perform partition-based graph abstraction (PAGA) analysis (Wolf et al., 2019), cellranger-derived count matrices were imported into scanpy (Wolf et al., 2018). The count matrix was normalized for size and log-transformed. High variable genes were selected with a minimum expression > 0.0125, a maximum expression < 3, and a minimum dispersion > 0.5. Twenty principal components were used to compute neighbors for UMAP. The leiden algorithm was used to perform clustering. PAGA analysis was performed using the default options in the scanpy.pl.paga function.

#### Bulk RNA-Seq experiments

Resultant 1×75 bp reads were demultiplexed and pseudoaligned with kallisto 0.42.5 software (https://pachterlab.github.io/kallisto/) to obtain tpm (transcripts per million) values (Bray et al., 2016). Pathway analyses were performed using GSEA software (Broad Institute) with the HALLMARK (Liberzon et al., 2015) reference. Normalized enrichment scores for individual pathways represent a comparison of the experimental timepoint for a given tissue fraction with the 0-d timepoint of the same tissue fraction (e.g., colon at exp d 9 vs. colon at exp d 0). SNEC pathway enrichment was compared against 0-d timepoints of both the distal colonic and anal tissue fractions. Global markers of SNEC were obtained by examining individual genes with expression ratio >4 compared to distal colonic and anal tissue fractions, merged across all times. Distinguishing pathways of marker genes were classified using functional annotation clustering on DAVID (https://david.ncifcrf.gov/) (Dennis et al., 2003).

#### Clonal analysis

The volume of Confetti+ clones was measured from DMI-reconstructed z-stacks of metaplasia. A region of interest corresponding to the xy-extent of the fluorescently marked cytosolic YFP+ or RFP+ cells was picked for each clone at each image plane in ImageJ. The volume of the clone was summed from each z plane and normalized to the total volume of the rete peg identified by its outlined structure from a reference nuclear stain. The allocation of clonal volume across tip, casing, or core compartments was made according to the following definitions: 1) tip - the single-cell thick periphery of the rete peg beginning at the abluminal vertex and extending until the peg has reached maximum cylindrical thickness, 2) casing - the single-cell thick peripheral layer of the cylindrical subportion of the rete peg, and 3) core - cells harboring a partially differentiated “spread” morphology in the interior of the rete peg. Sixty clones from combined 3-4 mice were measured per lineage tracing timepoint.

#### Statistical methods

Measurements were taken from distinct samples, as full-volume imaging allows the whole sample to be processed. Data with an exponential distribution were log-transformed to obtain normal distributions prior to statistical testing. Statistical comparisons between experimental and control conditions were performed using pairwise two-sided t-tests. Unless otherwise indicated, the Benjamini-Hochberg method of false discovery adjustment for multiple comparisons was applied to the p values. The adjusted p values (q values) are reported. Statistical tests were not performed on lineage tracing data when the effect size multiple was >3 or the number of samples was <3; in these cases the individual data points are plotted along with the mean.

#### Modeling

From the clonal data, we found that, over time, individual rete pegs in SNEC can become monoclonally labeled. In principle, such behavior could indicate that rete pegs are maintained by a single stem cell that renews through asymmetric division. Alternatively, such behavior could derive from an equipotent proliferative population that, like the normal intestinal colonic epithelium, undergoes neutral drift dynamics involving the chance replacement of neighboring tip cells lost through differentiation. To assess whether the tip proliferative cells of the rete pegs in SNEC represent an equipotent progenitor cell population, we turned to a statistical modeling-based approach.

Motivated by previous studies of crypt clone dynamics in the intestine (Lopez-Garcia et al., 2010) we considered a “voter model” to quantify cell dynamics within an equipotent progenitor population. In this formulation, a tip proliferative cell *S* carrying a hereditary label ‘a’ can replace its neighbor with label ‘b’ through symmetric cell duplication at a rate λ, described by the replacement “reaction:”

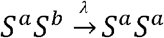

For an individual clone, the resulting pattern of lineage expansion and contraction follows a random walk or diffusion-like pattern. The mathematical formalism of this model has been detailed previously (Lopez-Garcia et al., 2010) and we only describe the essential findings here. When the renewing cells are arranged as a one-dimensional annulus or “ring,” such as that formed by intestinal stem cells at the crypt base, the smoothed distribution of clone sizes at early times, prior to significant clone fixation, takes the statistical scaling form:

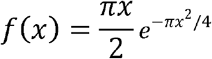

where 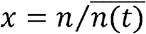 denotes the clone size *n* normalized by the average 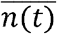. Therefore, a key test of whether the cellular system exhibits scaling is to plot the relationship 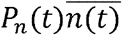 vs. 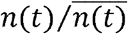, where *P* describes the probability of finding a clone of size *n* at time *t*. In a scaling system, clonal data from all timepoints should collapse onto the same time-independent scaling curve, *f*(*x*), provided the empirical tracing data are analyzed sufficiently early as to be within the scaling regime. Moreover, in the one-dimensional geometry, the surviving clone probability is given by:

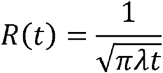

while the average size *n* varies as:

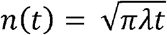

However, in the present case, the abluminal tip of the rete peg forms an ostensibly two-dimensional grid-like organization. In this case, the neutral drift dynamics are predicted to result in a scaling form in which the probability distribution converges over time to a simple exponential:

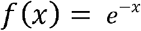

However, in stochastic simulations we found that in relatively small 2d compartments (i.e., lattices of 6×6 cells or smaller, within the size ranges of rete peg tip compartments), in which diagonal neighbor replacements are disallowed, the resulting clonal distributions could be reasonably modeled as 1d-type dynamics. Technically, an improved fit could be obtained with a linear combination of 1d (major contribution) and 2d (minor contribution) scaling functions (Figure S12a). However, for simplicity we primarily used the 1d voter model to study the tip clonal fate data. Importantly, when applied to the clonal data, we found that the 1d model provided a good fit to the data (Figure 7e).

To estimate the loss/replacement rate of the tip progenitors from the empirical longterm tracing of tip cells, we turned to the random walk equations that define exact solutions that account for clonal extinction and fixation within the scaling regime and during the later stages of clonal fixation. For the 1d voter model, the clone size probability distributions take the form:

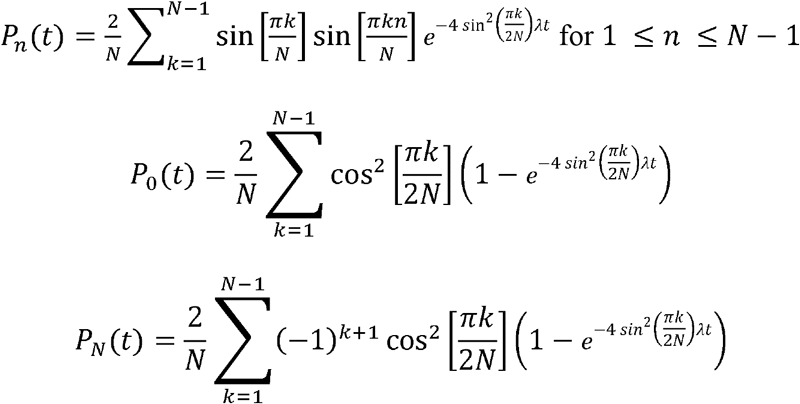

where *N* denotes the total size of the stem cell pool. To apply the model to the data, it is tempting to assume that the total number of renewing cells equates to the total number of tip proliferative cells. However, studies (Kozar et al., 2013) in the intestine have shown that the effective stem cell number *N* may be smaller than the number of cells that have renewal potential. Therefore, to address the experimental data, it makes sense to score clone size as a fraction of the total number of cells in the rete peg, defining the effective clone size as the fraction of the total area of the tip domain that are labeled, which translates to *n*/*N*. In this case, by fitting the distribution of persisting clone sizes 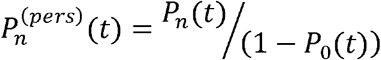 to the model prediction, we can estimate the effective rate:

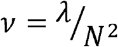

Moreover, to account for a practical delay between lineage tracing initiation (tamoxifen injection) and observation of clonal patches, the experimental data were shifted by *τ* = –1 week.

Since it was difficult to consistently resolve individual nuclei at the depth of the rete peg tip, we made use of clone morphologies in 3d images to quantify the fraction of labelled cells in rete pegs. The relationship of these fractions to cell numbers is simple if one assumes that tip compartment size is uniform. However, we found that the empirical fractional clone size converged to ~0.6 instead of the theoretical value of 1 (peg fixation), which challenged the fit at later tracing timepoints (> 12 wks, Figure S12b). We therefore devised a simple correction, based on our observation of heterogeneity in rete peg sizes. We found that rete peg diameters could range between 5-11 cells (data not shown). If one assumes that the tip cell compartment is of uniform absolute size, irrespective of the diameter of the rete peg, essentially implying that we have over-estimated the size of the tip compartment, then we can normalize the empirical clone size fraction by the maximally observed size. Such a normalization results in an improved fit (Figure 7f), with *ν* = 0.02 /wk, which is similar to the scaling kinetics found in small intestine and colon (Lopez-Garcia et al., 2010).

Finally, we note that the scaling behavior was robust to heterogeneity in tip compartment sizes. In particular, if *N* is not fixed, but is sampled from a statistical distribution, we could also generate good fits to the data. A key question is what is an appropriate distribution to use for values of *N*. A sample distribution can be obtained experimentally by examining the distribution of clone fractions at timepoints >12 wks, when the mean clone size does not change appreciably (Figure S12c). Thus, except for a small fraction of rete pegs that have become fully fixed as a result of a slow or infrequent transfer of cells from the tip to the casing compartments, this distribution provides insight into the spread of tip compartment sizes; that is, the tip compartment has been fixed. This tip “fixation distribution” can be sampled from a probability density function (Figure S12d) that has the empirical form:

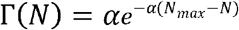

where N_max_ describes the maximum tip compartment size (we used N_max_ = 36 as it represents a 6×6 compartment). The result of this modification is a scaling system whose behavior represents the weighted sum of scaling subsystems evolving with different kinetics (λ/N^2^, assuming a fixed λ value). This model provided an excellent fit to the surviving clone size distribution, the growth in the mean clone size, and the shape of the clone size distribution throughout the tracing experiment (Figure S12e,f). However, it precludes reporting a single value of λ/N^2^. We note that previous theory has demonstrated that scaling behavior is preserved even with heterogeneity in values in λ/N^2^, provided that these values are sampled from a distribution with finite variance. Thus, any tip compartment size distribution meeting this criterion would result in scaling behavior.

## Table

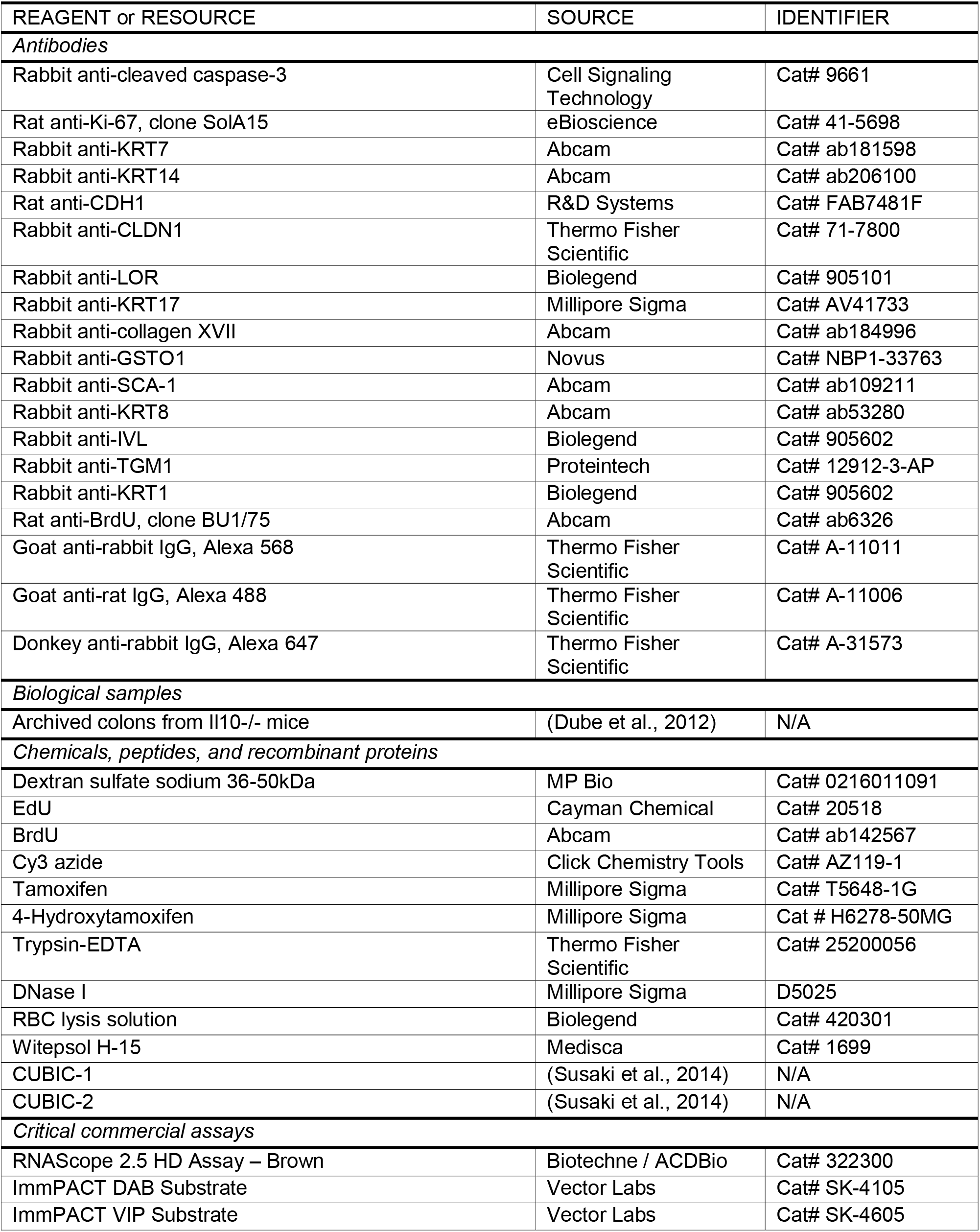
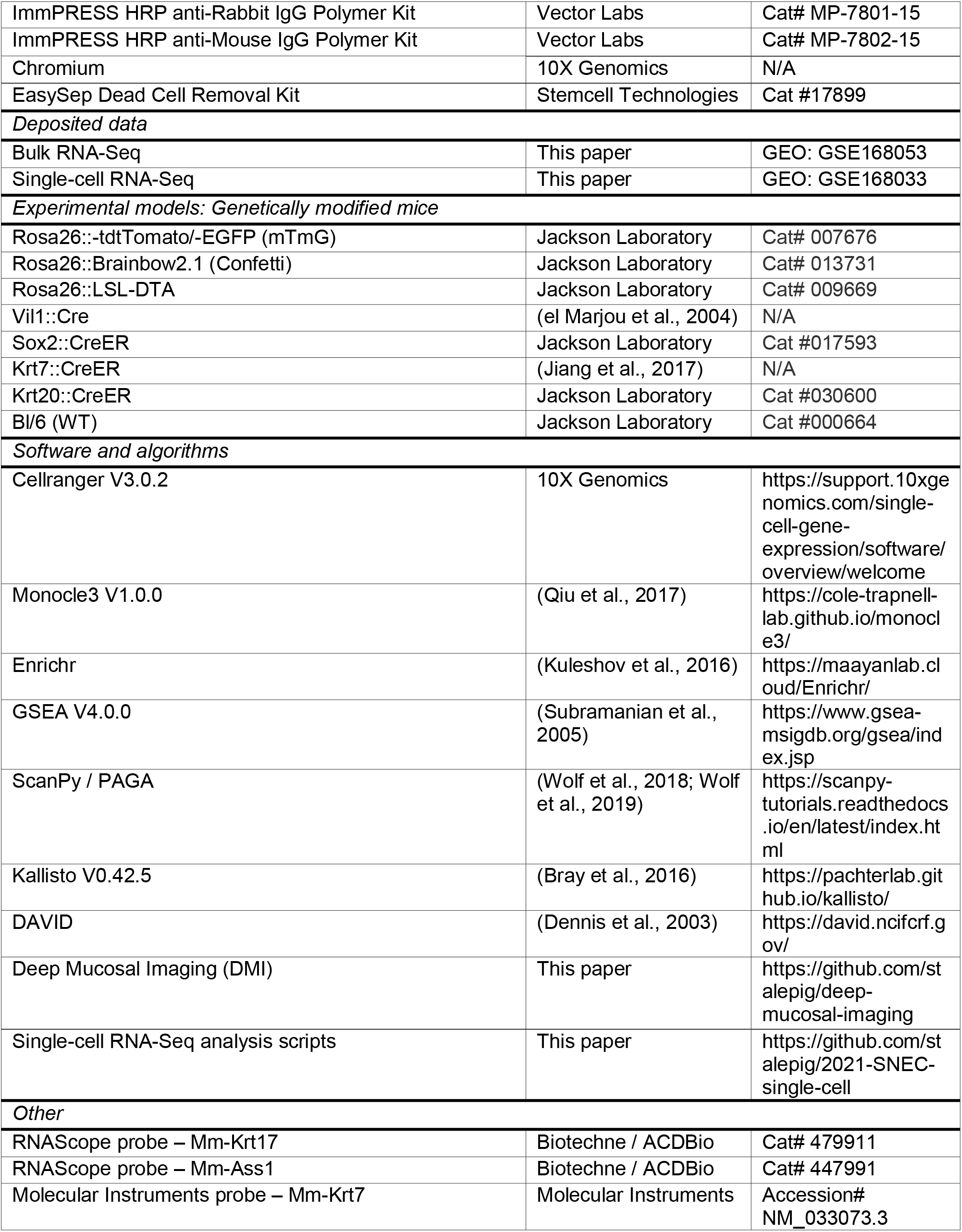

