## Supplementary material for "Anal skin-like epithelium mediates colonic wound healing": Figure S1

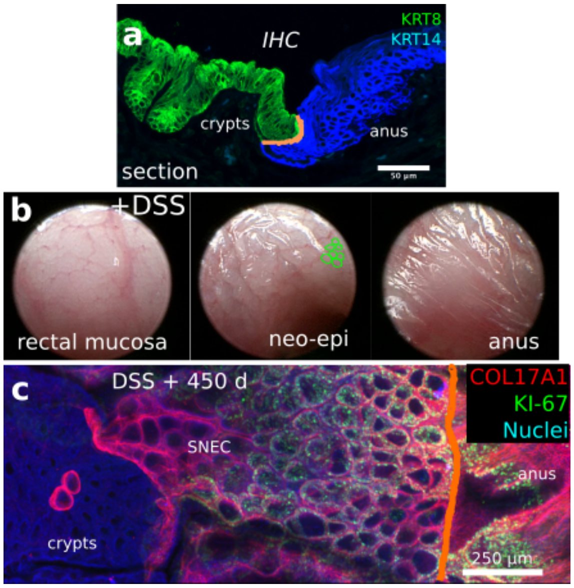


Squamous neo-epithelium of colon (SNEC) contributes to long-term colonic wound healing. Related to Figure 1. a) Thin-section photomicrograph of immunostained colonic (KRT8+) and anal squamous (KRT14+) epithelium reveals a sharp boundary between columnar and squamous epithelium at the dentate line (orange line), in the absence of injury. b) Mouse colonoscopy shows distinct structure of SNEC after DSS-induced injury. Example rete pegs are traced in green. c) In this *en face* projection of an interior section of a whole-mount staining, basal cells in SNEC express the squamous marker COL17A1 and the proliferation marker KI-67. Neighboring anal epithelium also expresses COL17A1 but colonic crypts do not. The dentate line is highlighted in orange. Error bars: SE.
