## Supplementary material for "Anal skin-like epithelium mediates colonic wound healing": Figure S2

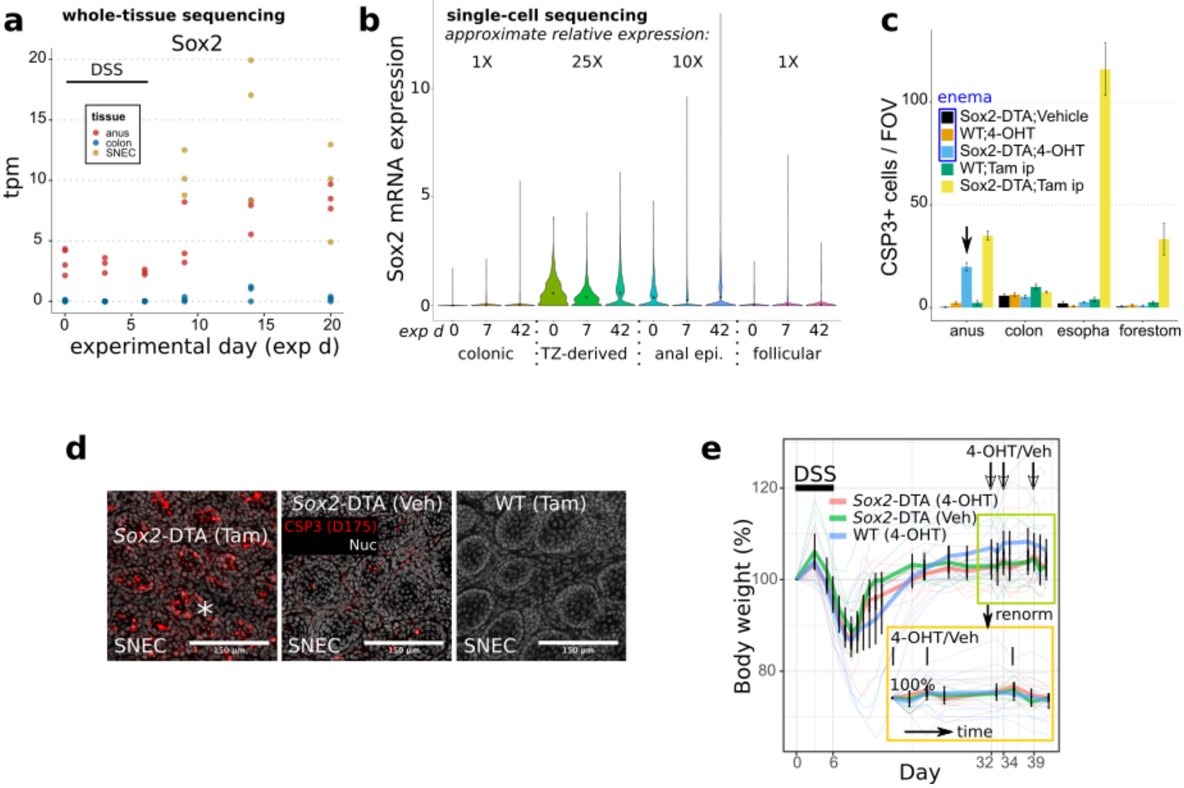


Induction of apoptosis in anorectal squamous cells with a conditional genetic construct. Related to Figure 1. a,b) Expression of Sox2 mRNA in whole-tissue sequencing (a) or single-cell sequencing (b) across different tissue or epithelial types shows consistent >10-fold enrichment for Sox2 expression in squamous epithelium vs. colonic epithelium. These sequencing data were obtained from experiments described in detail in Figures 4-6 and S2. c) We tested the specificity of targeting apoptotic signals to anal squamous tissue using Sox2-DTA mice (Sox2::CreER;Rosa26::LSL-DTA mice). Counting of caspase-3+ (CSP3, apoptotic) cells in response to enema (topical) or injection (systemic) of 4-OHT/tamoxifen shows specific apoptosis of anal epithelium after enema administration in Sox2-DTA mice (n=4 mice/condition). d) Whole-mount images demonstrate localization of cleaved caspase-3 (CSP3, asterisk) to SNEC cells after treatment of Sox2-DTA mice with DSS (to induce injury and SNEC formation) and 2 mg tamoxifen on exp d 32. e) In contrast to treatment during acute injury and repair, enema administration of 4-OHT after mucosal healing does not affect mouse body weight (n=6 mice/condition). Error bars: SE.
