## Supplementary material for "Anal skin-like epithelium mediates colonic wound healing": Figure S3

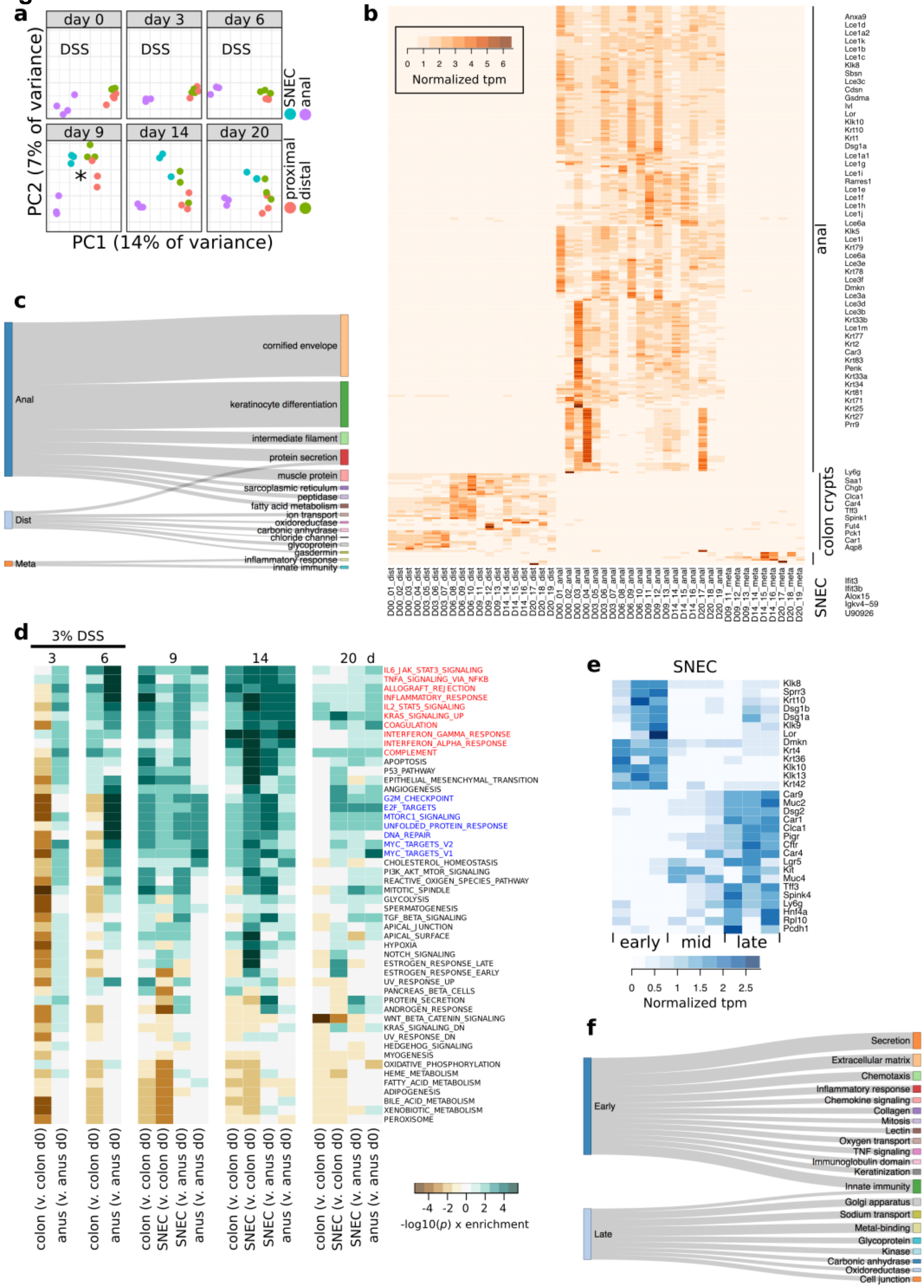

Whole-tissue profiling of colonic, anal, and SNEC samples. Related to Figure 1. a) Principal component plot of RNA-Seq data obtained from proximal-to-anal segments of colons from DSS-treated mice. Each dot represents a tissue sample. The initial emergence of SNEC is indicated by an asterisk (*). b-c) Tissue marker analysis (b) with gene ontology analysis (c) of anus, colonic crypts, or SNEC identify known squamous differentiation markers in anus, absorptive and secretory cell markers in colonic crypts, and interferon-inducible and inflammatory markers in SNEC ("meta"). d) GSEA of pathway activation/suppression at different timepoints versus the 0-d (uninjured) timepoint in colon or anus. Inflammatory pathways are highlighted in red. Proliferative pathways are highlighted in blue. "Enrichment" is defined as in the GSEA software. e-f) Changes in the overall expression characteristics of SNEC samples over time as shown at the single-gene level gene (e) and in ontology analysis (f). Early SNEC (d9) exhibits higher expression of inflammatory and squamous differentiation genes, while late SNEC (d20) shows elevated expression of colon-like genes. However, the absolute expression levels of these genes were ~10-fold less than in colon.
