## Supplementary material for "Anal skin-like epithelium mediates colonic wound healing": Figure S4

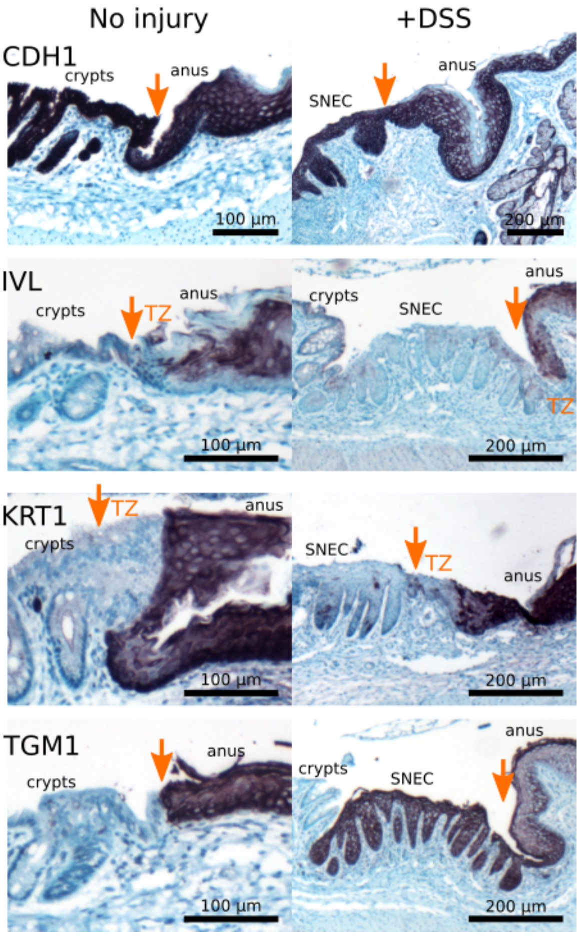


Related to Figure 2. Anal TZ represents a zone lacking terminal epidermal differentiation. Immunostaining of key markers of squamous differentiation at the anorectal junction, before (exp d 0) and after DSS (exp d 36) treatment. The orange arrow shows the dentate line, with the anal transition zone (TZ) marked immediately distal to it. Like SNEC, the TZ does not express IVL or KRT1 differentiation markers. Both colonic and anal epithelium express E-cadherin (CDH1). Anal epithelium, TZ, and SNEC express TGM1 (n=3 mice/stain).
