## Supplementary material for "Anal skin-like epithelium mediates colonic wound healing": Figure S5

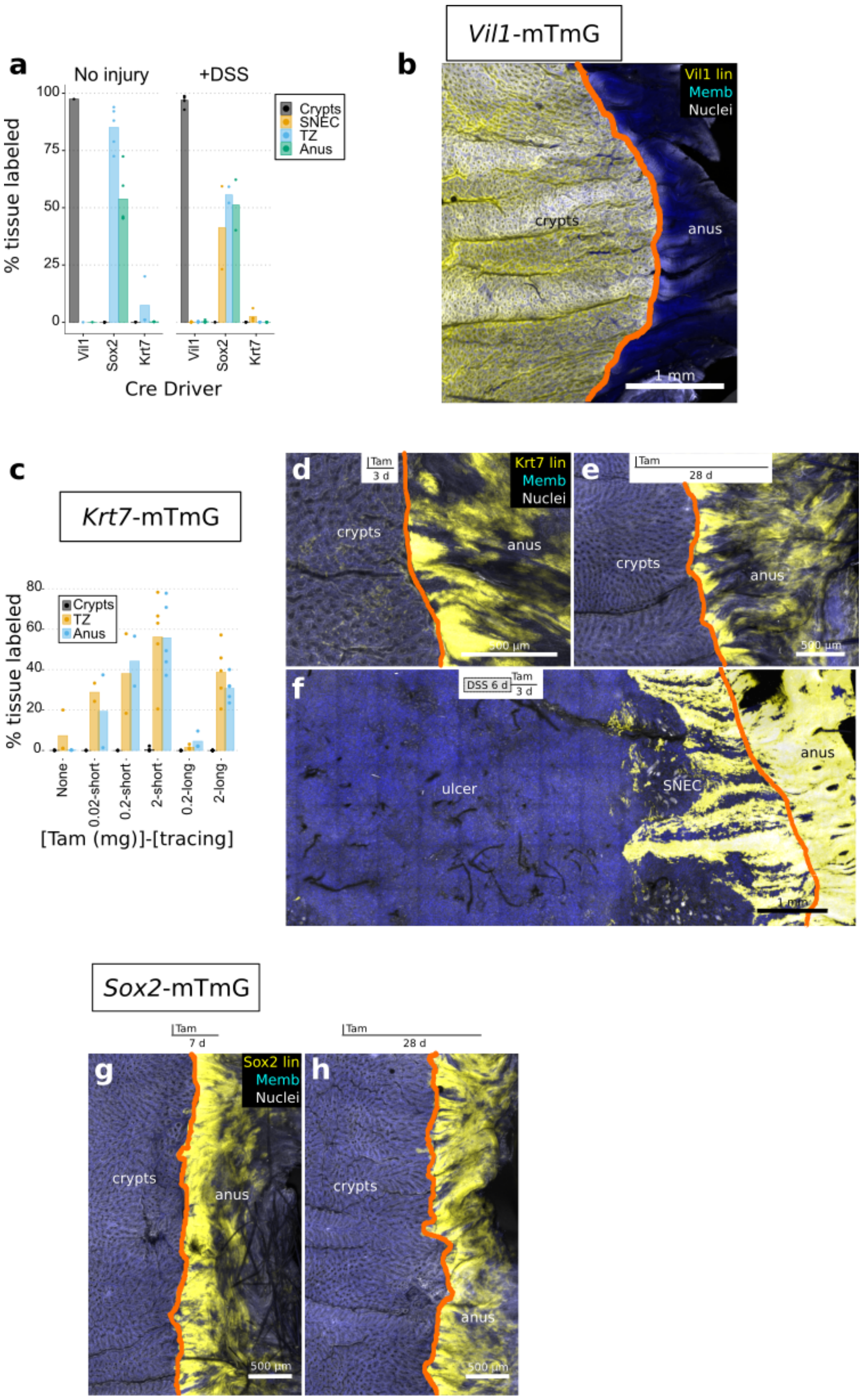


Assessing lineage contributions to the anorectal region in homeostasis and injury. Related to Figure 2. a) Quantitative tracing summary that shows the fraction of the epithelial surface labeled by various CreER driver mice in homeostasis and DSS-induced injury. Mice used: Vil1-mTmG (Vil1::Cre;Rosa26-mTmG, no tamoxifen injection), Sox2-mTmG (Sox2::CreER;Rosa26-mTmG, 2 mg i.p. tamoxifen), Krt7-mTmG (Krt7::CreER;Rosa26-mTmG, no tamoxifen injection). b) Whole-mount surface reconstruction of the anorectal region in Vil1-mTmG mice demonstrates specific labeling of crypts. The dentate line is marked in orange. c-f) Results from Krt7-mTmG mice. Doses of tamoxifen (0, 0.02, 0.2, or 2 mg) were administered to uninjured mice and the results imaged after different durations of tracing (3 d [short] or 28 d [long]). Quantification (c) shows the highest specificity of labeling for the anal TZ when only examining "leaky" recombination events that occur in the absence of tamoxifen. Higher doses of tamoxifen increased overall labeling but also labeled some squamous cells away from the TZ. Representative images after treatment with 2 mg tamoxifen in homeostasis are shown in d and e and after DSS-induced injury in f. g,h) Results from Sox2-mTmG mice showing that the homeostatic TZ remains labeled after both short (g) and long-term tracing (h).
