## Supplementary material for "Anal skin-like epithelium mediates colonic wound healing": Figure S6

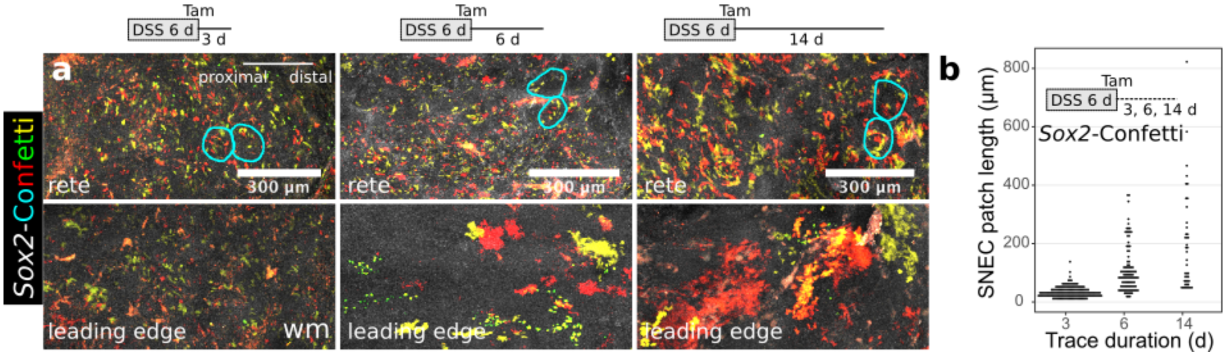


Sox2+ cells expand during acute colitis. Related to Figure 3. a) Short-term analysis in Sox2-Confetti mice during epithelial repair shows, in surface maximum intensity projections, the appearance of color patches after 3 d (exp d 9) and their expansion in the leading region (bottom row) after 6 (exp d 12) and 14 (exp d 20) d of tracing. Example rete pegs are outlined in light blue. b) Color patch size increases over time after DSS-induced injury. This shows that Sox2+ cells within SNEC drive the progressive re-epithelialization process.
