## Supplementary material for "Anal skin-like epithelium mediates colonic wound healing": Figure S7

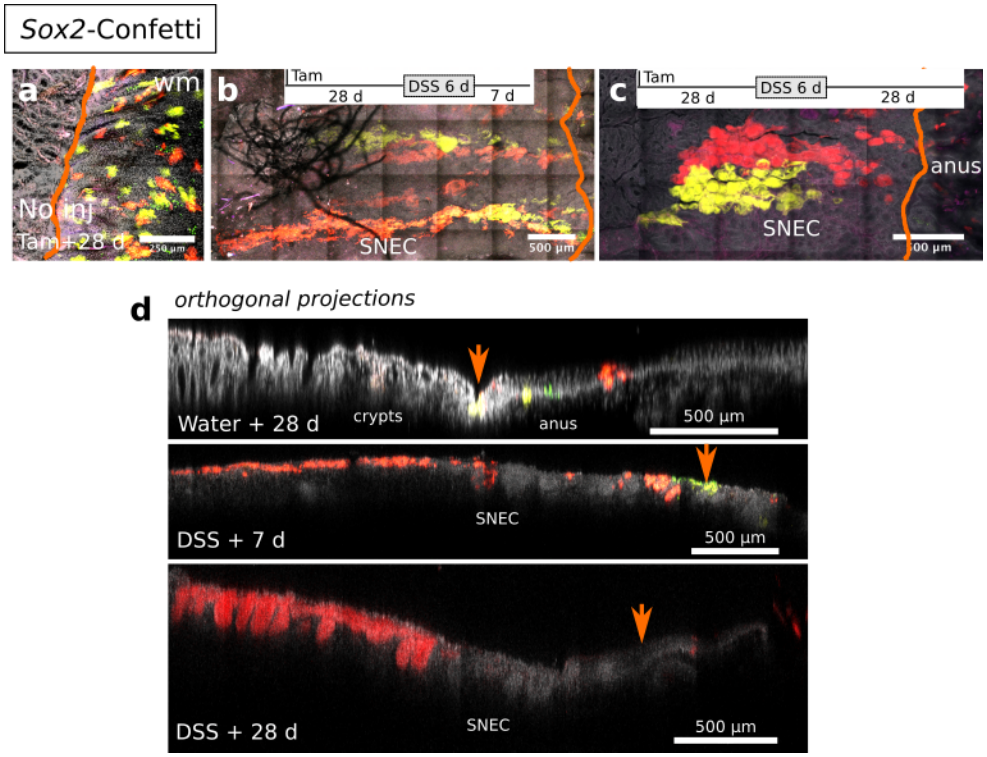


Emergence and remodeling of anal-derived cellular streams during SNEC formation. Related to Figure 3. a-c) Results from Sox2-Confetti mice show that clonal streams of cells forming SNEC arise from clonal collections of cells in the anus. The orange line marks the dentate line in whole-mount images. Surface reconstructions are displayed from uninjured colon (a) and from DSS-injured colon at 7 (b) and 28 d (c) after DSS withdrawal. d) Orthogonal projections from the images shown in a-c show the remodeling of the clonal streams from flat epithelium to rete peg epithelium. This demonstrates that SNEC is initially formed as a flat epithelium and that rete pegs emerge later. Orange arrows mark the dentate line.
