## Supplementary material for "Anal skin-like epithelium mediates colonic wound healing": Figure S8

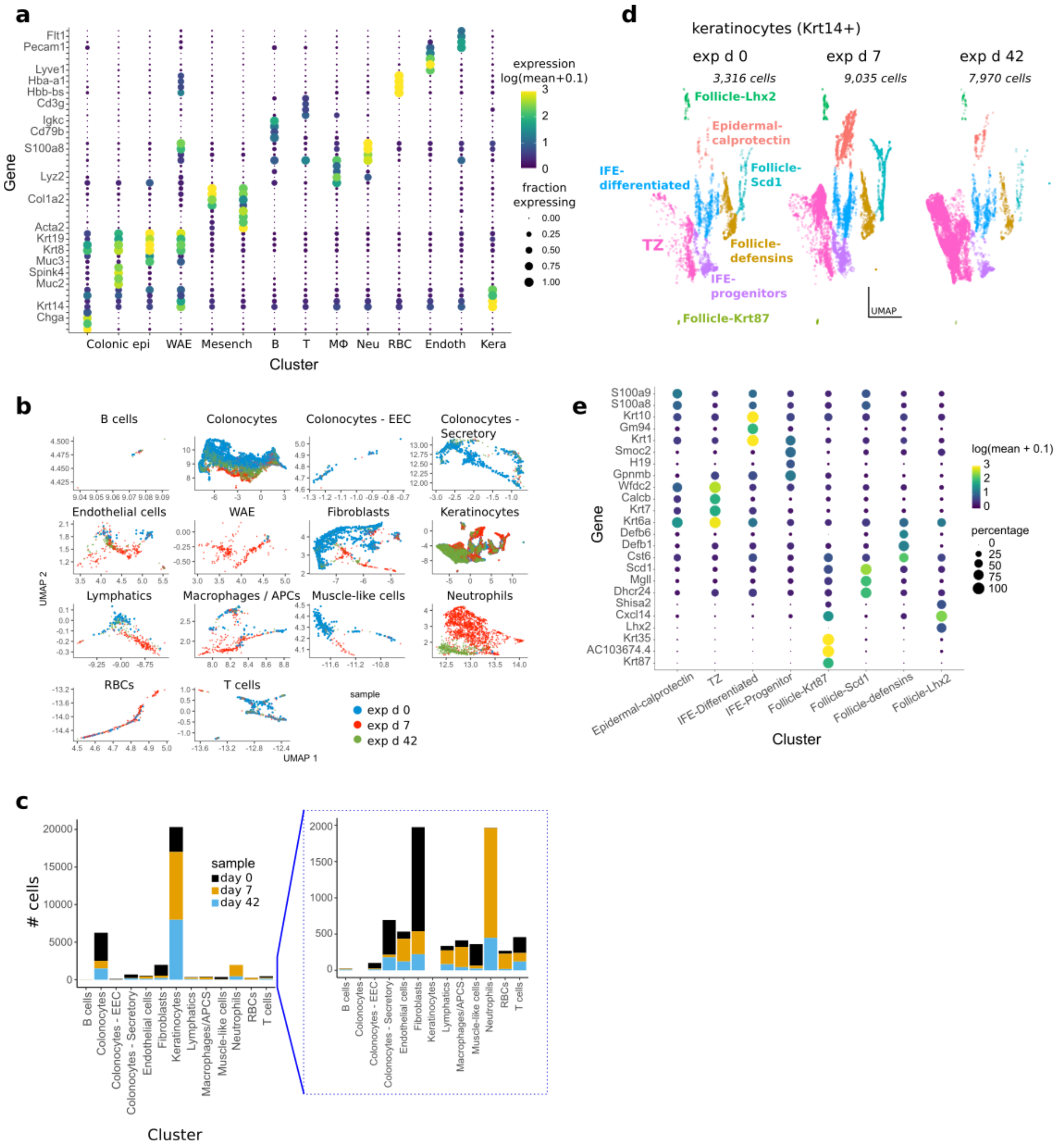


Transcriptomic characterization of cell type-specific responses during DSS colitis. Related to Figure 4. a) The dotted plot shows major gene markers driving the top-level clustering of the merged scRNA-Seq data obtained from the anorectal region at exp d 0, 7, and 42. Abbreviations: epi = epithelium, WAE = wound associated epithelium, Mesench = mesenchyme, B = B cells, T = T cells, Mϕ = macrophages, Neu = neutrophils, RBC = red blood cells, Endoth = endothelium, Kera = keratinocytes. b,c) UMAP visualization of changes in individual cell clusters (b) and their abundance (c) associated with DSS-induced injury; clusters where the dot colors do not overlap represent cell types undergoing broad shifts in transcriptional profile. d) The time-resolved UMAP visualization shows major subtypes of keratinocytes. Note the increased abundance of TZ-related cells at exp d 42, which corresponds to the presence of SNEC. IFE = interfollicular epithelium. e) The dotted plot reveals markers used to classify keratinocytes shown in d.
