## Supplementary material for "Anal skin-like epithelium mediates colonic wound healing": Figure S9

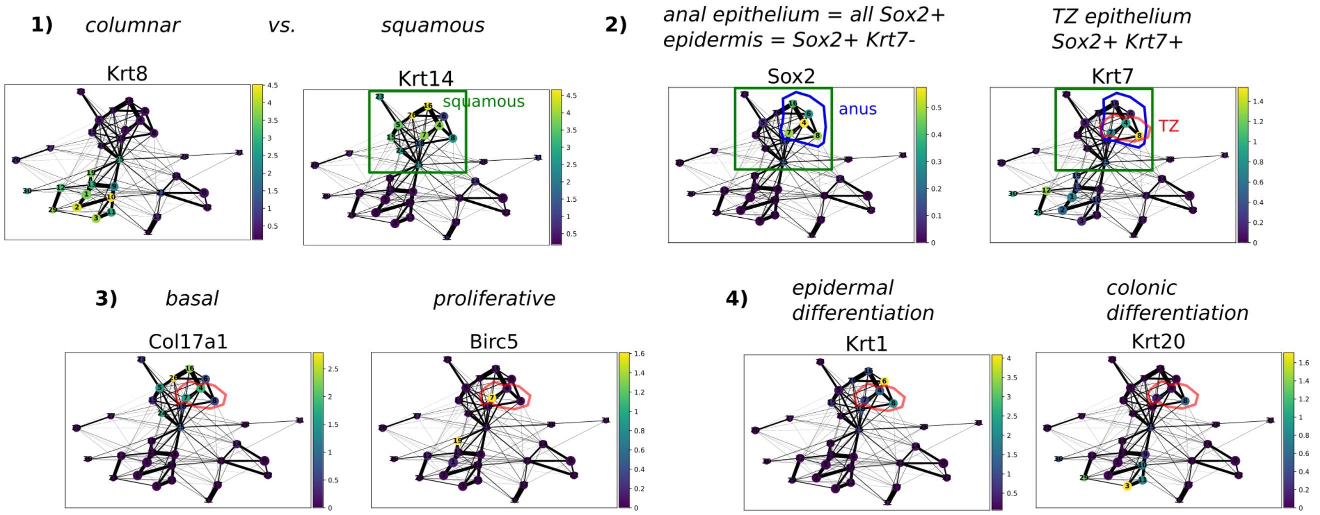


Graph analysis of single-cell mucosal transcriptomes obtained from exp d 0. Related to Figure 4. The count matrix was imported into scanpy, size-normalized and log-normalized, trimmed to highly variable genes, dimensionally reduced using 20 principal components, and projected using UMAP. Clusters were identified using the leiden algorithm. Similarity relationships between clusters were computed using partition-based graph abstraction (PAGA). Each point on the plot represents a cluster of cells. The color shading indicates the expression level of the gene per cluster. A "gating" strategy was applied such that squamous cells (Krt14+) were distinguished from colonocytes (Krt8+). Keratinocytes could be divided into anal squamous epithelium (Sox2+) or follicular-type epithelium (Sox2-negative). Within Sox2+ epithelium, three clusters of $Krt7^{hi}$ cells were identified; these clusters putatively represent the anal TZ. Similar to the analytical pipeline performed in monocle3, two of the TZ clusters represented basal cells (Col17a1+), with one cluster harboring highly proliferative cells (Birc5+). The suprabasal TZ cluster was positive for the colonocyte marker Krt20 but lacked high expression of the squamous cell differentiation marker Krt1. Note that the TZ clusters are embedded within the squamous cell graph. This suggests that TZ cells are fundamentally squamous in nature, but have elevated expression of colonic markers. This finding is in agreement with the results independently obtained in Figure 4.
