## Supplementary material for "Anal skin-like epithelium mediates colonic wound healing": Figure S10

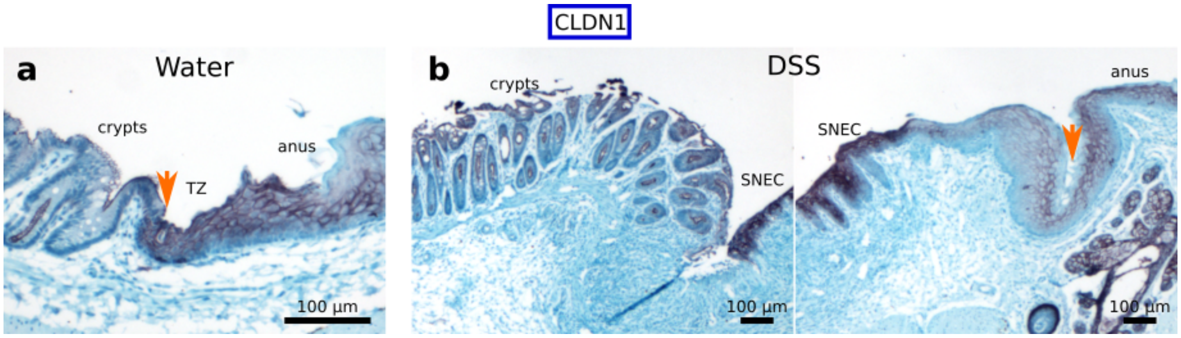


SNEC cells exhibit elevated expression of claudin-1. Related to Figure 6. a) Immunohistochemical labeling of CLDN1 at the uninjured anorectal junction shows staining in rectal crypts and the TZ. b) Staining after DSS-induced injury shows elevated expression of CLDN1 protein in SNEC relative to anal epidermis and the original TZ. Orange arrows mark the dentate line.
