## Supplementary material for "Anal skin-like epithelium mediates colonic wound healing": Figure S11

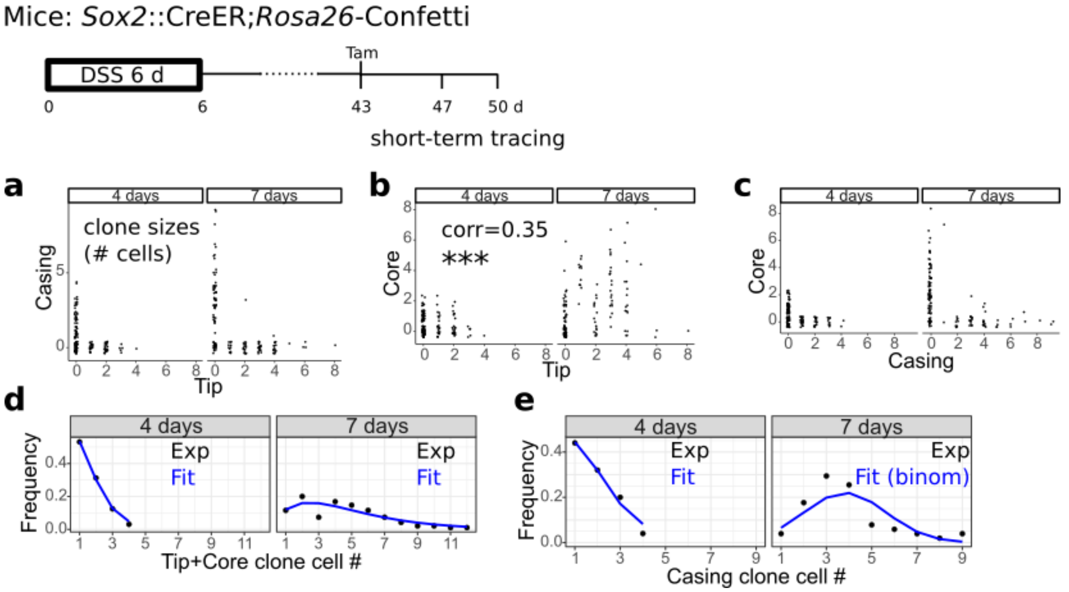


Short-term lineage tracing of progenitor cell populations in SNEC. Related to Figure 7. a-e) Short-term lineage traces (4 and 7 days, 0.5 mg tamoxifen per mouse, n=3 mice/timepoint) in SNEC begun 43 d after DSS treatment in Sox2-Confetti mice. The correlations between the number of cells in each color patch in the tip, casing, and core compartments are plotted (a-c). Only tip and core cell occupancy were correlated (corr., Pearson’s r), supporting the regeneration of core cells by tip cells and separate function of the casing domain. The size distributions of tip+core (d) and casing (e) cells were plotted and show smooth distributions that are fit by exponential or binomial functions.
