## Supplementary material for "Anal skin-like epithelium mediates colonic wound healing": Figure S12

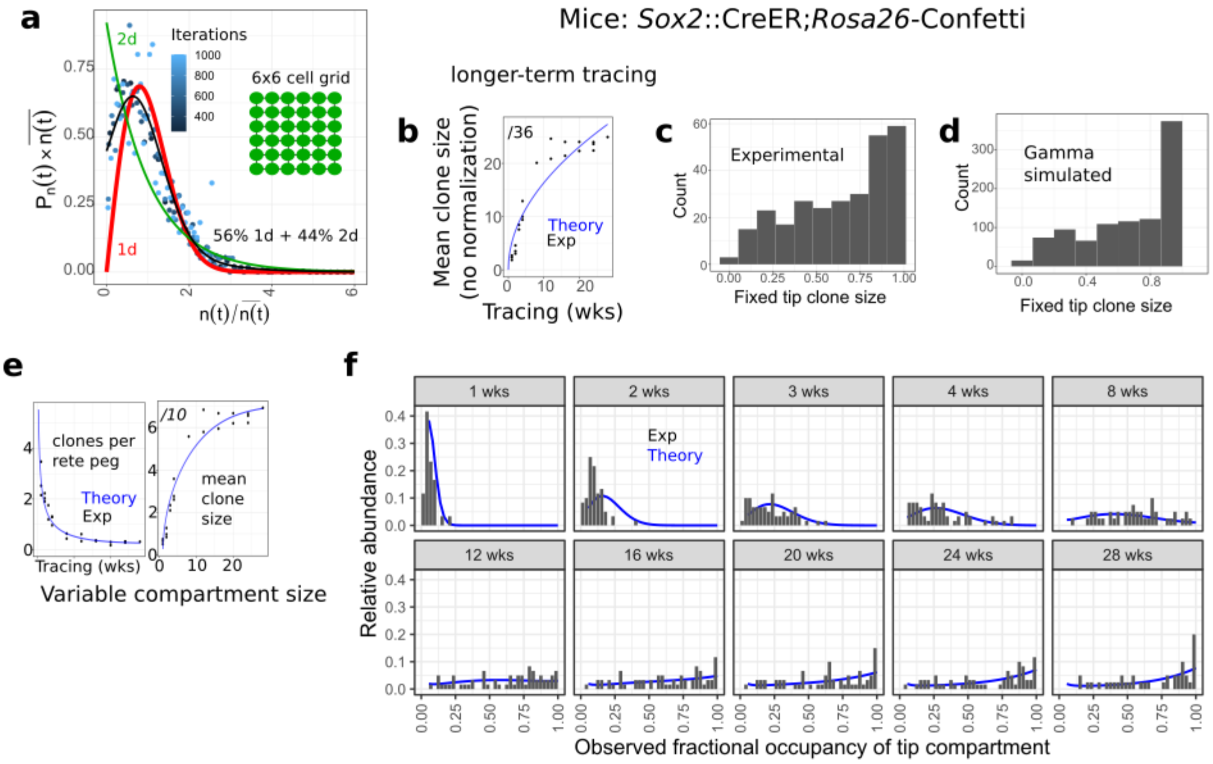


Modeling long-term regeneration in SNEC with variable tip compartment sizing. Related to Figure 7. a) Simulation of clone sizes emerging at different times (iterations) from random replacement within an equipotent 6x6 square grid of cells; the time-invariant scaling form is shown (see "Methods"). Diagonal replacement is not allowed in the simulation. A 1d scaling function provided a good fit to the simulated clone sizes. b) Fitting of the theoretical evolution ($\lambda/N^2$=0.008 /wk) of the un-normalized mean clone size over longer-term (1-28 wks) tracing. The fitting at later timepoints (>12 wks) is suboptimal because the mean clone fractional size converges to ~0.6 (<1). c) Shown is the distribution of empirical tip clone sizes expressed as a fraction of the total tip domain identified from imaging. The distribution is aggregated from data after 12 wks of tracing. This "fixation distribution" explains why the mean clone size asymptotically approaches 0.6 within the timeframe of the tracing experiment. d) The fixation distribution can be modeled with a simulated gamma function. e,f) Adjusting the neutral drift model to account for heterogeneity in tip compartment sizes provides a good fit to the empirical elimination and growth rate of clones (e), as well as the empirical shape of the tip clonal size distribution (f).
